# Sequestration of growth cone surface proteins by cytoplasmic Lrrtm2 induces *de novo* amygdala innervation by cerebral cortex associative neurons

**DOI:** 10.64898/2026.04.08.716720

**Authors:** Dustin E. Tillman, Omer Durak, Priya Veeraraghavan, John E. Froberg, Garrett Wheeler, Bogdan Budnik, Jeffrey D. Macklis

## Abstract

Precise establishment of distinct cerebral cortex circuits is essential for sensorimotor function, high-level cognition, and cross-modality integration and association. Although an increasing set of molecular controls over subtype-specific cortical wiring have been identified, much less is known about how molecules in growth cones (GCs) regulate precise long-range projection of axons through complex environments, or how dysregulation of GC molecular machinery disrupts precision of circuit formation. Here, we discover a generalizable mechanism for regulation of precise circuit wiring by focusing on callosal projection neurons (CPN), which link cortical hemispheres via the corpus callosum. CPN are centrally involved in associative and cognitive function, and are often disrupted in people with autism spectrum disorders (ASD) and intellectual disabilities (ID). We identify dysregulated subcellular CPN GC proteomes *in vivo* after CPN-specific deletion of *Bcl11a*/*Ctip1*, a transcription factor (TF) with variants that cause ASD/ID in humans, and validate localization of dysregulated proteins to CPN GCs *ex vivo*. We identify that disruption of Lrrtm2 – a canonically postsynaptic transmembrane protein – in CPN GCs specifically induces *de novo* innervation of the amygdala, an evolutionarily ancient regulator of social behavior, cognition, and anxiety that is abnormally activated in humans with ASD. Mechanistically, we identify that deletion of *Bcl11a* from CPN disrupts targeting of Lrrtm2 to CPN GC membranes, causing cytoplasmic sequestration of key CPN GC surface proteins, and resulting in aberrant innervation of basolateral amygdala (BLA), which is typically targeted by evolutionarily older archicortex. Together, this work connects deletion of a causal ASD/ID TF, dysregulation of a non-canonical control over GC surface protein remodeling, and formation of a *de novo*, subtype-specific circuit between cerebral cortex and BLA – similar mechanisms likely generalize across neuron subtypes. These results expand conceptual understanding of how diverse circuits are precisely constructed and potentially evolve, and how coordinated dysregulation of GC molecules can disrupt precise subtype-specific circuitry, contributing to diverse neurodevelopmental and neuropsychiatric disorders.

## Introduction

Cerebral cortex is unique to mammals, and it has increased in size across evolution to enhance control over integrated sensation, skilled movement, and high-level cognition^1^. This expansion is closely linked with diversification of cortical projection neuron subtypes, which precisely innervate distant targets to form functionally distinct circuits^2^. Prior studies have identified an increasing set of combinatorial molecular controls in projection neuron cell bodies that regulate subtype specification and circuit wiring^3^. However, much less is known about downstream processes by which these diverse nuclear molecules effect subtype-specific axon guidance and precise circuit formation *in vivo*.

GCs are specialized subcellular compartments at tips of growing axons that rapidly integrate extracellular signals to control axon guidance, then mature into presynapses^4^. The immense distance separating GCs from their parent somata (10^3^ – 10^5^ cell body diameters), in addition to substantial emerging evidence, indicate that GCs mostly function semi-autonomously via local synthesis and degradation of GC proteins^5–9^. Elegant *in vitro* work has deeply investigated these processes, and characterized molecular pathways that are activated by binding of specific extracellular cues at distinct transmembrane receptors, ultimately resulting in cytoskeletal remodeling that regulates the direction of axon extension for precise innervation of distinct target areas^10^.

We previously developed experimental and analytical approaches to purify GCs from specific neuron subtypes in developing mouse brains, enabling in *vivo* identification of their transcripts and proteins^11,12^. We have employed these approaches to identify that GC transcriptomes are subtype-specific^13^, dysregulated after deletion of corticospinal TF *Bcl11b*/*Ctip2*^14^, localized via distinct sequence features^15^, and enable precise wiring via homophilic interactions^16^. This work highlights the power of subcellular investigation to directly discover local controls over circuit development, function, and maintenance, and elucidate how subcellular disruptions in subtype-specific neurons drive disease.

CPN are an exemplar projection neuron subtype that function in sensorimotor, associative, integrative, cognitive, and behavioral circuits by projecting to precise homotopic targets in layer II/II of contralateral cortex via the corpus callosum to distribute information between key functional areas of the two hemispheres^17^. Many neurodevelopmental and neurological disorders are linked with disrupted CPN circuitry, including ASD/ID^18^, callosum agenesis^19^, and split-brain syndrome^20^. Intriguingly, ASD-risk genes are enriched in human CPN and other superficial layer projection neuron subtypes^21,22^, and perturbing human ASD-risk genes often alters CPN circuits^23–25^. Further, long-range projections are disrupted in humans with ASD^26^, as is precise synchronicity between the cortical hemispheres^27^. Together, these results underscore the close association of CPN with typical high-level cognition, and their functional implication in ASD/ID, highlighting the importance of investigating precise CPN circuit formation to better understand complex cognitive, behavioral, and related neurodevelopmental disorders.

Recently, we investigated developmental defects and causal molecular mechanisms of dysfunctional *Bcl11a*^28^, a TF that controls acquisition of CPN identity, precision of CPN targeting, radial positioning of CPN, and organization of cortical sensory areas^29–31^. Notably, humans and mice with *BCL11A* microdeletion or loss-of-function mutations display key clinical features of ASD, ID, and developmental delay^32–36^, leading to its classification as a high-confidence, syndromic, fully penetrant ASD/ID-risk gene by the Simons Foundation Autism Research Initiative (SFARI)^37^.

In this recent work, we identify that CPN-specific *Bcl11a* deletion in mice leads to defects in social novelty preference and working memory, and three primary CPN circuit abnormalities^28^: 1) promiscuous, inaccurate contralateral targeting; 2) rerouting via the evolutionarily older anterior commissure; and, most strikingly, 3) aberrant, *de novo* innervation of BLA, a regulator of social behavior, cognition, and anxiety that is abnormally activated in humans with ASD^38–40^. Mechanistically, we also elucidate transcriptomic dysregulation of *Bcl11a*^-/-^ GCs via both disrupted expression and trafficking, and identify that two dysregulated CPN GC RNAs, *Mmp24* and *Pcdhαc2*, function downstream of *Bcl11a* to regulate precise contralateral innervation^28^. However, neither *Mmp24* nor *Pcdhαc2* drive *de novo* BLA innervation, suggesting that additional GC molecular regulators, likely direct GC protein effectors, might function downstream of *Bcl11a* to produce this strikingly aberrant CPN circuit.

The semi-autonomy of GCs motivates investigation of subcellular proteomes to elucidate controls over subtype-specific circuit development and implementation. Although accessibility of low-input RNA-seq has led to use of RNA abundance as a proxy for protein levels (at steady state for most cells, ∼50% of protein variance is due to variance in RNA^41,42^), this correlation weakens substantially in cells that are transitioning between states or responding to stimuli^43,44^, and is 2-fold weaker in neurons at baseline^45,46^. GCs navigate through diverse extracellular environments, locally synthesize and/or degrade proteins in response to distinct guidance cues, and contain many proteins that are synthesized in somata, then efficiently trafficked to GCs for localized function^47–49^. Thus, GC RNA abundance is a much less reliable indicator of GC protein levels; our prior work in CPN GCs identified an R^2^ of 0.05 between GC RNA and GC protein abundances^11^. Therefore, direct investigation of GC proteomes is highly motivated toward discovery of unidentified regulators controlling key, unexplored aspects of normal and aberrant subtype-specific circuit formation.

Here, we identify dysregulated proteomes of *Bcl11a*^-/-^ CPN GCs by enhancing approaches for *in vivo* ultra-low-input subcellular proteomics. We validate presence of select dysregulated proteins in CPN GCs *ex vivo*, and elucidate striking function of one exemplar candidate: Lrrtm2. We identify that Lrrtm2, known canonically as a postsynaptic transmembrane protein, is localized in developing CPN GCs, and misprocessed after *Bcl11a* deletion. Physiological, low-level overexpression (OE) of cytoplasmic Lrrtm2 in CPN, but not membrane-inserted Lrrtm2 or its ligand (Nrxn), causes *de novo* innervation of the BLA by CPN. Mechanistic investigations identify that this non-canonical cytoplasmic Lrrtm2 binds to many GC membrane proteins that function in axon guidance (e.g. Nrxn), sequestering them from CPN GC surfaces, and resulting in aberrant BLA targeting. These results elucidate non-canonical GC regulators controlling subtype-specific circuit wiring during *in vivo* development, with direct implications for ASD/ID. Similar non-canonical mechanisms likely generalize to other neuron subtypes, and demonstrate the discovery power of investigating subtype-specific subcellular proteomes in developing neurons.

## Results

### Subcellular proteomes of *Bcl11a*^**-/-**^ CPN are dysregulated

Our recent work elucidated subcellular dysregulation of *Bcl11a*^-/-^ CPN transcriptomes via both abnormal trafficking and transcription (and without changes in CPN somata for many RNAs), and identified two dysregulated GC RNAs that function in CPN contralateral targeting^28^. However, neither of these GC RNAs contributes to the strikingly aberrant, *de novo* innervation of BLA by *Bcl11a*^-/-^ CPN. These results motivated direct subcellular investigation of GC proteins that potentially cause formation of this aberrant circuitry.

We aimed to identify GC proteins regulating this striking circuit abnormality via enhanced approaches for ultra-low-input subcellular proteomics (see Materials and Methods). Briefly, CPN of *Bcl11a*^fl/fl^ mice were fluorescently labeled (± Cre) via *in utero* electroporation (IUE) at embryonic day (E) 14.5. At postnatal day (P) 3, CPN somata were purified by microdissection, trituration, and fluorescence-activated cell sorting (FACS), and CPN GCs were purified in parallel via microdissection, subcellular fractionation, and fluorescent small particle sorting (FSPS) of bulk growth cone fraction (GCF) input^11,12^ (Fig. 1a; Supp. Fig. 1a, b). Purified CPN somata and GCs were analyzed via independent runs of tandem mass tag mass spectrometry (TMT MS), utilizing advanced ultra-low-input proteomic approaches that we and others have recently developed^50–53^ (Supp. Tables 1-3). Critically, RNA and protein protection assays identify that CPN GCs rapidly seal, remaining intact and membrane-enclosed (Supp. Fig. 1c, d). Pilot subcellular proteomes reveal a ∼5-fold increase in proteomic sensitivity relative to our earlier GC work via improved TMT MS methods^11^ (Supp. Fig. 1e, f). In parallel, *Bcl11a*^+/+^ CPN GCs were also purified at P0 and P3 to enable further investigation of GC proteins potentially regulating axon extension rates and midline crossing, both perturbed in *Bcl11a*^-/-^ CPN^28^ (Supp. Fig. 1g-i).

**Figure 1:**
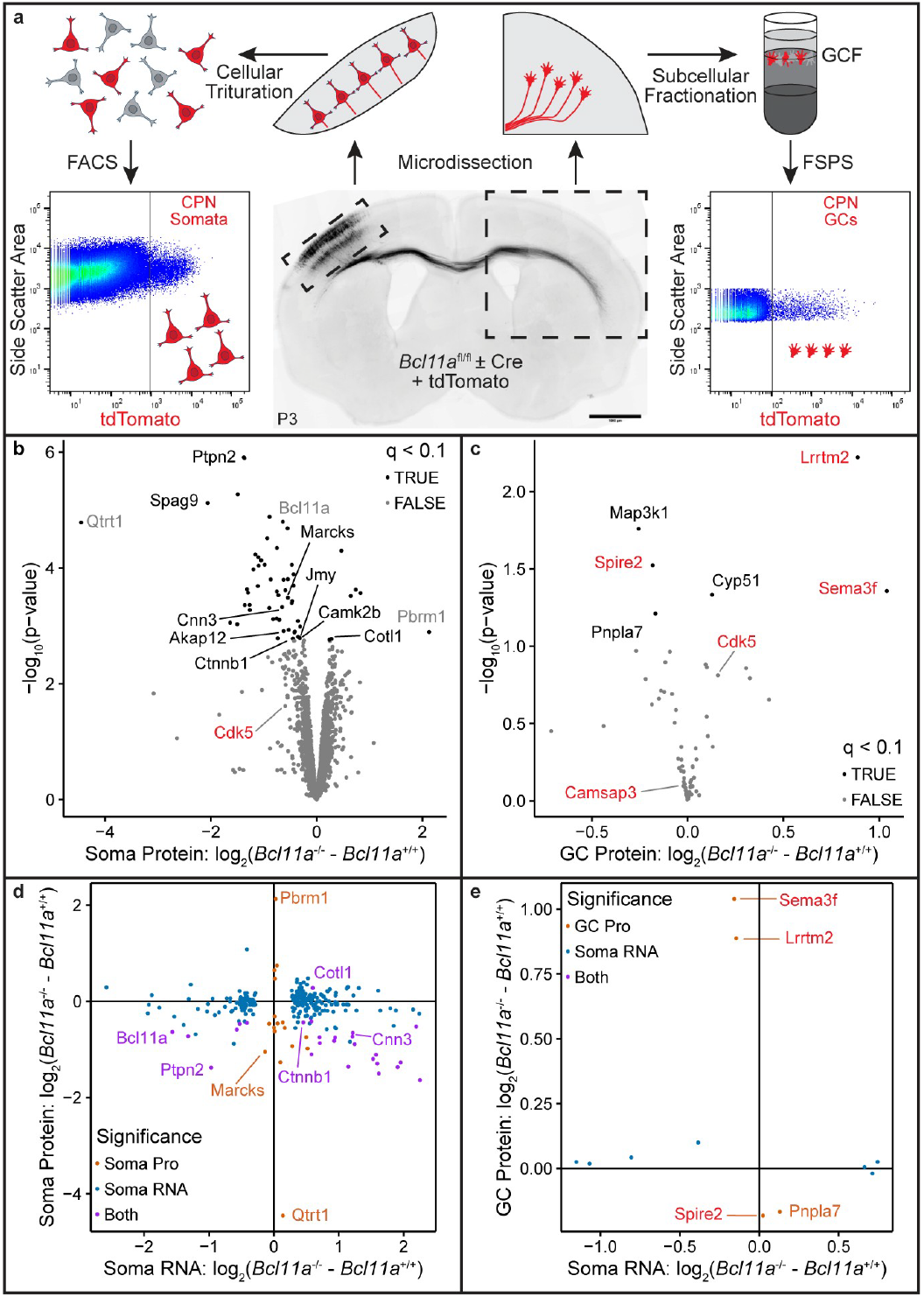
Subcellular proteomics identifies candidate GC proteins regulating CPN circuit formation. **(a)** CPN of Bcl11a^fl/fl^ mice are fluorescently labeled (± Cre) via E14.5 IUE. At P3, somata are purified via microdissection, cell trituration, and FACS, and GCs are purified via microdissection, subcellular fractionation, and FSPS. **(b, c)** Proteins with differential abundance in Bcl11a^-/-^ CPN somata **(b)** or GCs **(c)**. Candidate protein names are red, GC protein names are black, and other notable protein names are gray. **(d, e)** Genes with differential RNA abundance in Bcl11a^-/-^ CPN somata or differential protein abundance in Bcl11a^-/-^ CPN somata **(d)** or GCs **(e)**. Candidate protein names are red. Soma RNA data from^26^. q < 0.1 for all tests. Scale bars: 1000 um.

To identify GC proteins that potentially control CPN circuit formation downstream of *Bcl11a*, and with the BLA in particular, we evaluated proteomes across subcellular compartments and *Bcl11a* genotypes. Consistent with GC semi-autonomy and localized protein function, proteomes of *Bcl11a*^+/+^ and *Bcl11a*^-/-^ somata form distinct clusters, and GC proteomes cluster by condition (Supp. Fig. 1j, k). While only 15% of differentially abundant proteins in *Bcl11a*^-/-^ somata (q < 0.1) are known to function in GCs (Fig. 1b), all six differentially abundant proteins in *Bcl11a*^-/-^ GCs (q < 0.1) participate in GC molecular networks, highlighting the power of directly investigating GCs to identify subcellular dysregulation of key GC proteins. The previously described functions of these six dysregulated GC proteins include synaptogenesis (Lrrtm2^54,55^), axon pathfinding (Sema3f^56^), actin nucleation (Spire2^57^), signal transduction (Map3k1^58^, Cdk5^59^ (q = 0.15), and lipid alteration (Pnpla7^60^, Cyp51^61^).

Intriguingly, integration of previous CPN subcellular transcriptomes^13,28^ identifies that 73% of molecules with significant changes in RNA and protein levels upon *Bcl11a* deletion (q < 0.1) have a negative correlation between RNA and protein abundances (Fig. 1e), suggesting disrupted regulation over subcellular protein localization after *Bcl11a* deletion, potentially via altered translation and/or axonal transport^15,62^. Further, and quite strikingly, none of the six proteins with dysregulated GC levels upon *Bcl11a* deletion have a significant change in cell body RNA abundance after *Bcl11a* deletion (Fig. 1f). This result, consistent with our recent RNA work^11–14,28^, demonstrates the power of *in vivo* ultra-low-input GC proteomics to identify proteins directly controlling circuit formation that are undetected by more standard RNA-seq and/or cell body proteomic approaches.

We filtered differentially abundant GC proteins based on previously described functions, focusing on molecules that are known to either regulate axon growth or guidance, or function in synapses. We reasoned that these GC proteins might be most likely to regulate GC function, and contribute to circuit disruptions of *Bcl11a*^-/-^ CPN. Using this strategy, we identified Lrrtm2 (Leucine-rich repeat transmembrane neuronal protein 2) and Sema3f (Semaphorin-3f) as top-tier candidates for validation and functional investigation as potential exemplar regulators of CPN circuit development.

Lrrtm2 is canonically known as a postsynaptic ligand for key Neurexins (Nrxns) that organize presynapses once they mature from GCs^54,55,63–65^, and Sema3f is known to function as a secreted semaphorin that controls axon targeting in other circuits via binding to Neuropilin-2 (Nrp2) and/or Plexin-A3 (PlexA3) receptors^56,66–69^. Spire2 (less abundant in *Bcl11a*^-/-^ GCs) and Camsap3 (decreased abundance in P0 *Bcl11a*^+/+^ GCs) were selected as second-tier candidates, due to their functions in actin nucleation at tips of growing protrusions^57^, and microtubule organization during neuron polarization^70–74^, respectively. Cdk5 is (non-significantly) more abundant in *Bcl11a*^-/-^ GCs (q = 0.15), so we also aimed to validate presence of this protein in GCs, because of its many neuronal functions, including in GC motility^59,75,76^. The rich literature describing functions of these GC proteins in other circuits, or more broadly across neuronal and cell biology, nominated them as candidates to function locally in CPN GCs for regulation of precise circuit formation *in vivo*.

### Candidate GC proteins are localized to CPN GCs *in vivo*

To verify identification of candidate GC proteins by ultra-low-input mass spectrometry, we aimed to validate *in vivo* localization of candidate proteins in CPN GCs. As a first assessment, we investigated enrichment of candidates in bulk cortical GCs via western blot of the GCF relative to the post-nuclear homogenate (PNH) of cortex (Fig. 2a), as we and others have done^11–13^. As expected, Gap43 is enriched in the GCF, and Map2 is depleted from the GCF (Fig. 2b; Supp. Fig. 2a), validating the accuracy of this approach. Interestingly, via this gross metric of non-subtype-specific GCF, ligands of Lrrtm2 and Sema3f (Nrxn and Nrp2/PlexA3, respectively), but not either candidate GC protein itself, are enriched in the GCF, indicating high cortical abundance of Lrrtm2 and Sema3f, and localization of their receptors to bulk cortical GCs. Spire2, Camsap3, and Cdk5 are also not enriched in the broad, non-subtype-specific GCF by western blot, suggesting high cortical abundance, and not excluding presence or enrichment in subtype-specific CPN GCs.

**Figure 2:**
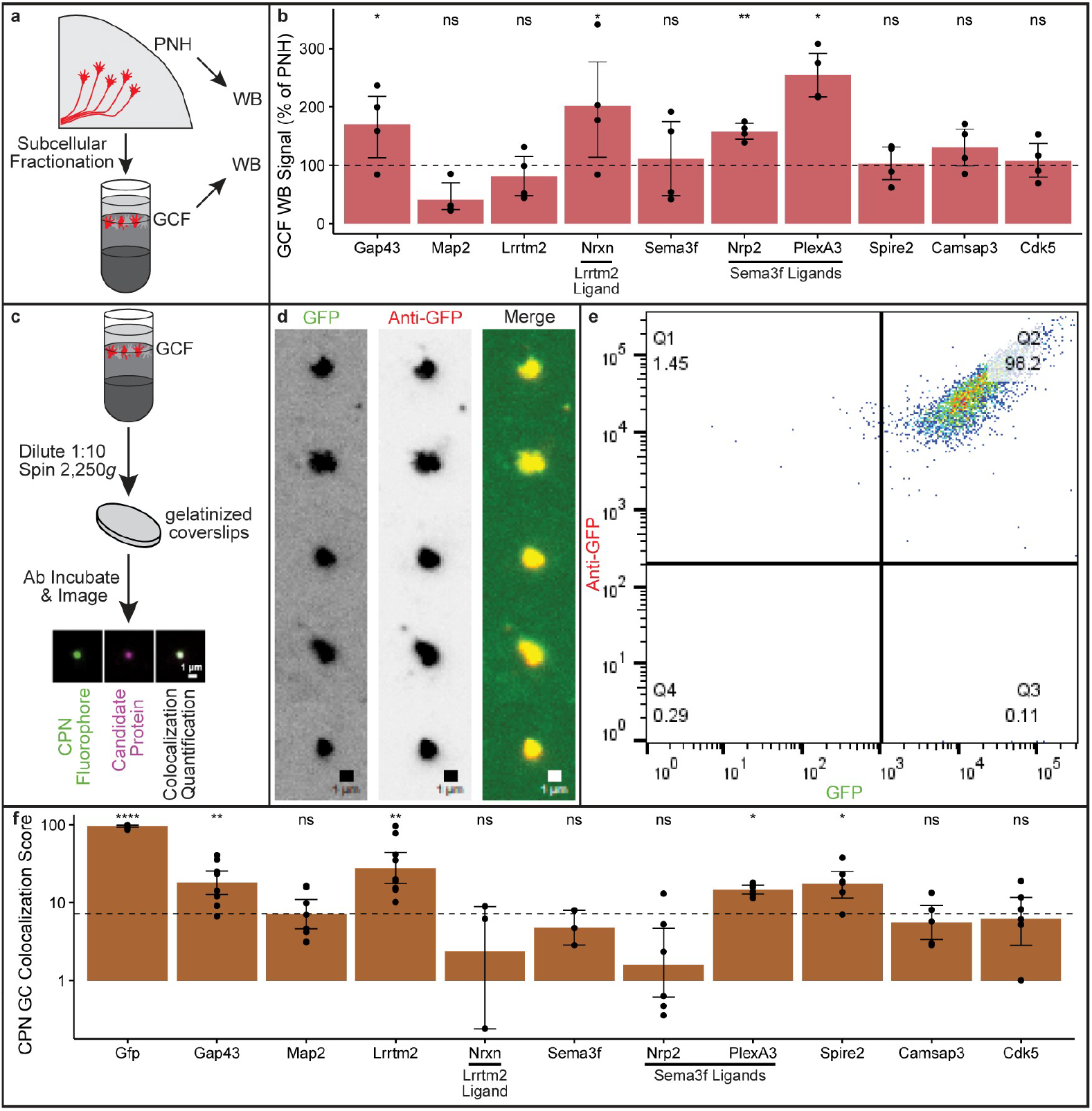
Lrrtm2 and Sema3f are highly localized to CPN GCs *in vivo*. **(a)** Subcellular fractionation of cortical PNH yields GCF for western blot analysis. **(b)** Western blot validates abundance of candidate proteins in bulk cortical GCs relative to cortex. Dotted line is no enrichment in GCF. **(c)** Centrifuging GCF onto gelatin-coated coverslips adheres GCs, enabling *ex vivo* colocalization analysis of GC candidates with fluorescent CPN GCs. **(d)** Representative CPN GCs (green) labeled with anti-GFP (red). **(e)** Representative colocalization plot of CPN GCs (green) labeled with anti-GFP (red). **(f)** Candidate protein CPN colocalization scores. Dotted line is mean CPN colocalization score of Map2. Error bars: 95% CI. Statistics: one-sided Student’s t-test (****: p < 0.0001, ***: p < 0.001; **: p < 0.01; *: p < 0.05; ns: p > 0.05). Scale bars: 1 um.

To more deeply investigate subtype-specific localization of select candidate proteins to CPN GCs *in vivo*, we adapted approaches previously reported for immunofluorescence of adherent synaptosomes^77^. Briefly, we centrifuged dilute GCF containing CPN GCs that were fluorescently labeled via E14.5 IUE, fixed adherent P3 GCs, incubated with antibodies against candidate proteins, and quantified a “colocalization score” that equally weighs abundance in, and specificity for, CPN GCs (Fig. 2c). As expected, GFP antibodies colocalize with native GFP-labeled CPN GCs (Fig. 2d, e), demonstrating how this approach accurately identifies colocalization of candidate proteins in subtype-specific GCs *ex vivo*. Similarly, Gap43 is more colocalized than Map2 (Fig. 2f; Supp. Fig. 2b), although the compressed nature of these GC colocalization scores suggest a limited dynamic range of this approach, potentially due to steric competition for limited antibody binding sites on small (∼300-500 nm diameter^11^) CPN GCs.

Importantly, colocalization of candidate proteins in CPN GCs (Fig. 2f) indicates that subtype-specific validation of GC localization is critical, especially for low-abundance neuron subtypes. Cruder western blot assessment of the cortical GCF only partially captures results from the more refined assay of subtype-specific, adherent GCs. Based on the high colocalization of Lrrtm2, PlexA3 (a Sema3f receptor), and Spire2 with CPN GCs *ex vivo*, and considering previously described functions of these proteins in synaptogenesis, axon guidance, and actin growth, respectively, we focused functional investigations on Lrrtm2 and Sema3f, as well as their associated receptors, to identify potential functions of these proteins in regulating construction of CPN circuitry.

### Lrrtm2 regulates aberrant amygdala innervation by CPN

Lrrtm2 is a leucine-rich repeat transmembrane protein that others have identified on postsynaptic membranes, where it binds presynaptic Nrxn to control synapse formation and modulate circuit activity^54,55,63,78–80^. More broadly, other leucine-rich repeat proteins (e.g. Slit), including some with transmembrane helices (e.g. Trk, Linx/Islr2, Lumican), are known to control axon guidance cell-autonomously and/or non-cell-autonomously across other circuits^81–84^. However, no Lrrtm proteins have been previously reported to function at earlier developmental stages, or in GCs, which mature into presynapses after innervating target areas.

Since Lrrtm2 is significantly more abundant in *Bcl11a*^-/-^ CPN GCs, we investigated potential function(s) of Lrrtm2 in CPN circuit wiring by transfecting CPN with a low-level Lrrtm2 OE construct via E14.5 IUE, and analyzing CPN circuits at P7 and P21. To distinguish between cell-autonomous and non-cell-autonomous Lrrtm2 functions, we also used an OE plasmid for Nrxn3aMS4, which lacks exon 4, enabling it to bind Lrrtm2^54,63,64^.

Notably, RNAs that are less abundant in *Bcl11a*^-/-^ CPN cell bodies^28^ are enriched for gene ontology (GO) terms related to membrane protein localization and signal peptide processing (Supp. Fig. 3a). Therefore, we hypothesized that the Lrrtm2 signal peptide, conserved for leucine-rich repeat proteins^85^, might be improperly processed in *Bcl11a*^-/-^ CPN, leading to a select increase in abundance of non-canonical cytoplasmic, not standard membrane-inserted, Lrrtm2 in *Bcl11a*^-/-^ CPN GCs.

To test this hypothesis, we engineered a construct for low-level OE of Lrrtm2 without its native signal sequence (cytLrrtm2) for comparison with a construct for low-level OE of membrane-inserted Lrrtm2 (memLrrtm2). Critically, transfection of HEK cells with these constructs results in localization of memLrrtm2 to membrane (Supp. Fig. 3b), and localization of cytLrrtm2 to cytoplasm (Supp. Fig. 3c), relative to a cytoplasmic control – both constructs increase Lrrtm2 abundance by ∼3.5-fold in HEK cells (Supp. Fig. 3d).

Strikingly, low-level OE of non-canonical cytoplasmic Lrrtm2 (cytLrrtm2^OE^) in CPN causes specific innervation of BLA relative to a tdTomato control (Fig. 3a), reproducing the *de novo* circuit abnormality identified in *Bcl11a*^-/-^ CPN^28^. In contrast, low-level OE of canonical membrane-inserted Lrrtm2 (memLrrtm2^OE^) or its ligand (Nrxn3aMS4^OE^) in CPN does not result in BLA innervation, revealing that Lrrtm2 must be atypically localized in cytoplasm, not membrane, to cell-autonomously cause aberrant CPN innervation of BLA.

**Figure 3:**
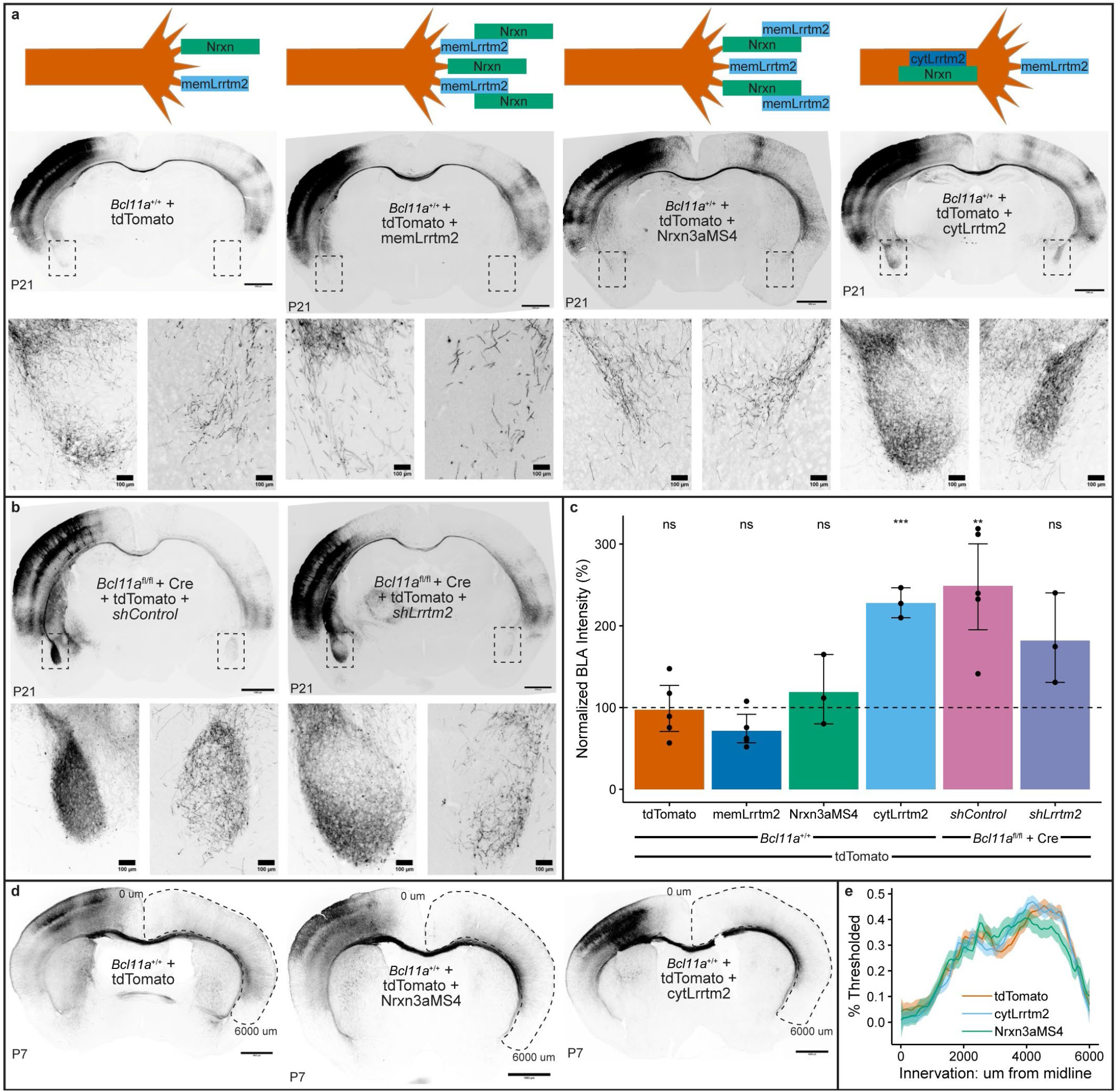
Cytoplasmic Lrrtm2 controls aberrant BLA innervation by CPN. **(a, b)** CPN are transfected via E14.5 IUE with plasmids encoding for low-level overexpression of candidate proteins **(a)** or shRNAs **(b)** and imaged at P21. Schematics define how low-level overexpression might impact abundance and localization of select GC proteins, and insets highlight BLA. **(c)** Quantification of BLA innervation by CPN across conditions. Dotted line is innervation by *Bcl11a*^+/+^ CPN. **(d)** CPN circuits across conditions at P7. **(e)** Quantification of contralateral precision by CPN across conditions. Error bars: 95% CI. Statistics: two-sided Student’s t-test (***: p < 0.001; **: p < 0.01; ns: p > 0.05). Scale bars: 1000 um. Inset scale bars: 1 um.

To validate this identified, non-canonical function of Lrrtm2 in CPN circuit formation, we performed a “rescue” experiment by transfecting *Bcl11a*^-/-^ CPN with shRNAs against either a non-mammalian RNA sequence (*shControl*) or Lrrtm2 (*shLrrtm2*) via E14.5 IUE. This partial Lrrtm2 KD (∼50% in P1 CPN somata; Supp. Fig. 3e) partially rescues BLA innervation by *Bcl11a*^-/-^ CPN (Fig. 3b, c), confirming that Lrrtm2 contributes to BLA targeting of *Bcl11a*^-/-^ CPN. Importantly, IUE sites are replicable (Supp. Fig. 3f, g), demonstrating that this striking circuit phenotype is not a result of differential CPN positioning following E14.5 IUE.

Intriguingly, cytLrrtm2^OE^ CPN still precisely innervate contralateral cortex (Fig. 3d, e), even though this aspect of circuit formation is substantially perturbed with *Bcl11a*^-/-^ CPN^28^. Further, investigation of Sema3f and its receptors (Nrp2 and PlexA3), known to control axon guidance, radial migration, and/or extension speed in other circuits^29,56,66– 69,86–89^, identifies a subtle role of this molecular network in CPN axon extension speed that is independent of its control over cell body migration (Supp. Fig. 4-6; detection of this subtle phenotype required a recently developed mosaic genetic analysis system^90^). Together, these results indicate separable regulation over distinct aspects of circuit wiring by distinct molecular controls, and demonstrate how broad disruptions to RNA expression and subcellular localization in *Bcl11a*^-/-^ CPN lead to separable and select circuit defects.

Thus, for mechanistic investigations, we focused on how non-canonical cytoplasmic Lrrtm2 disrupts normal circuit wiring and causes aberrant innervation of the BLA by CPN.

### cytLrrtm2 sequesters Nrxn from CPN GC surfaces *in vivo*

Since cytLrrtm2^OE^, but not memLrrtm2^OE^ or Nrxn3aMS4^OE^, leads to BLA innervation by CPN, and since RNAs encoding for membrane protein localization and signal processing GO terms are dysregulated in *Bcl11a*^-/-^ CPN, we hypothesized that *Bcl11a* deletion increases abundance of misprocessed, cytoplasmic Lrrtm2 in CPN GCs, leading to sequestration of axon guidance receptors directly or indirectly, and resulting in aberrant CPN circuitry. To investigate subtype-specific abundances of proteins on CPN GC surfaces, we employed surface-labeling FSPS (slFSPS), an approach that our lab recently developed^14^. Briefly, GCF with labeled CPN GCs via E14.5 IUE are incubated with antibodies against a candidate protein of interest, and unbound antibodies are removed by centrifugation prior to FSPS analysis (Fig. 4a). We validated this approach by verifying that GPI-GFP, but not cytGFP, is identified on GC surfaces by slFSPS of WT CPN (Fig. 4b)^14^.

**Figure 4:**
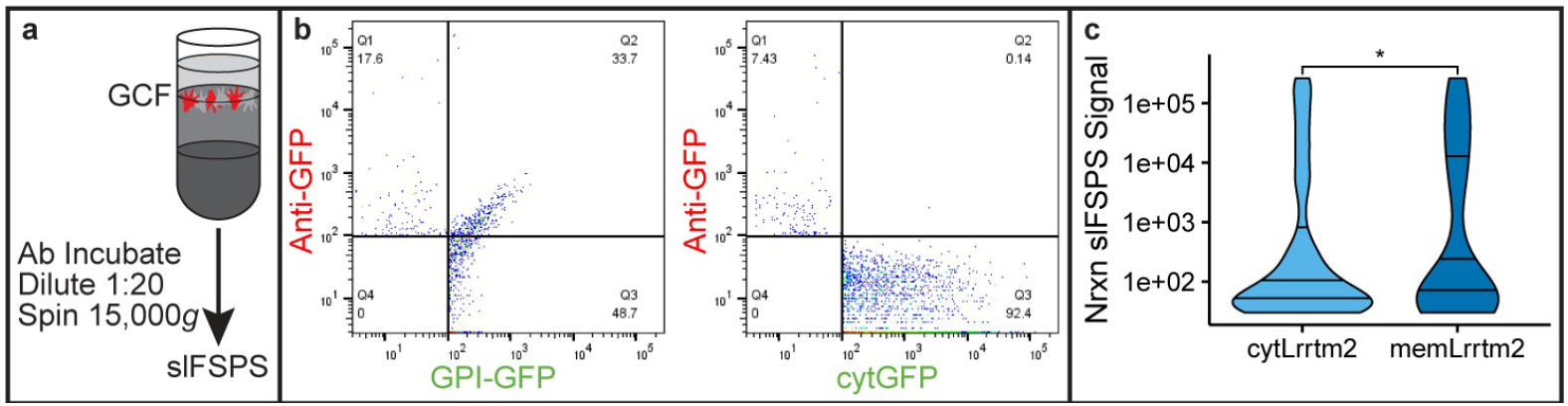
Nrxn is sequestered from CPN GC surfaces by cytoplasmic Lrrtm2. **(a)** Incubating GCF with antibodies enables quantification of GC surface protein abundance via slFSPS. **(b)** Representative slFSPS GFP plots for GPI-GFP and cytGFP. **(c)** Quantification of Nrxn abundance on CPN GC surfaces across conditions. Solid lines: 25^th^, 50^th^, and 75^th^ percentiles. Statistics: two-sided Student’s t-test (*: p < 0.05).

To first investigate whether Nrxn, an Lrrtm2 ligand that functions in axon guidance^91–94^, might be sequestered from GC surfaces by cytoplasmic Lrrtm2, we transfected CPN with either cytLrrtm2^OE^ or memLrrtm2^OE^ constructs. Strikingly, Nrxn is significantly less abundant on GC surfaces in cytLrrtm2^OE^ CPN relative to memLrrtm2^OE^ CPN (Fig. 4c; Supp. Fig. 7a, b). This reveals that cytLrrtm2 sequesters Nrxn from CPN GC surfaces, potentially contributing to aberrant BLA targeting by cytLrrtm2^OE^ CPN. To investigate whether Nrxn is directly bound by cytLrrtm2, and to investigate whether cytLrrtm2 might sequester additional proteins, we investigated other potential binding partners of cytLrrtm2 *in silico* and *in vivo*.

### cytLrrtm2 binds Nrxn and other GC guidance molecules

Although direct interactions between Lrrtm2 and Nrxn are well-described^54,63,64^, we hypothesized that cytLrrtm2 may either indirectly decrease abundance of Nrxn on CPN GC surfaces, and/or bind other key GC proteins, resulting in aberrant innervation of the BLA by CPN. To first elucidate the range of potential Lrrtm2 interactors, we performed an *in silico* immunoprecipitation (IP) of Lrrtm2 against mouse proteome with AlphaPulldown^95,96^, which uses AlphaFold-Multimer to predict interactions with “bait” proteins^97,98^ (Fig. 5a). Confirming known direct interactions, this *in silico* analysis identifies Lrrtm2-Nrxn complexes (Fig. 5b; Supp. Fig. 7c; Supp. Table 4), further supporting the likelihood that cytLrrtm2 may directly bind Nrxn in CPN GCs. Notably, Lrrtm2 is also predicted by this analysis to interact with other GC adhesion proteins, including Pcdh15, Epha4, and Cdh7 (Fig. 5c; Supp. Fig. 7d, h, i, m, n). All of these molecules are known to regulate axon guidance in other neuron subtypes^99–102^, suggesting that cytLrrtm2 might bind and inhibit multiple proteins in parallel to dysregulate *in vivo* CPN circuit formation, acting as a “dominant negative”.

**Figure 5:**
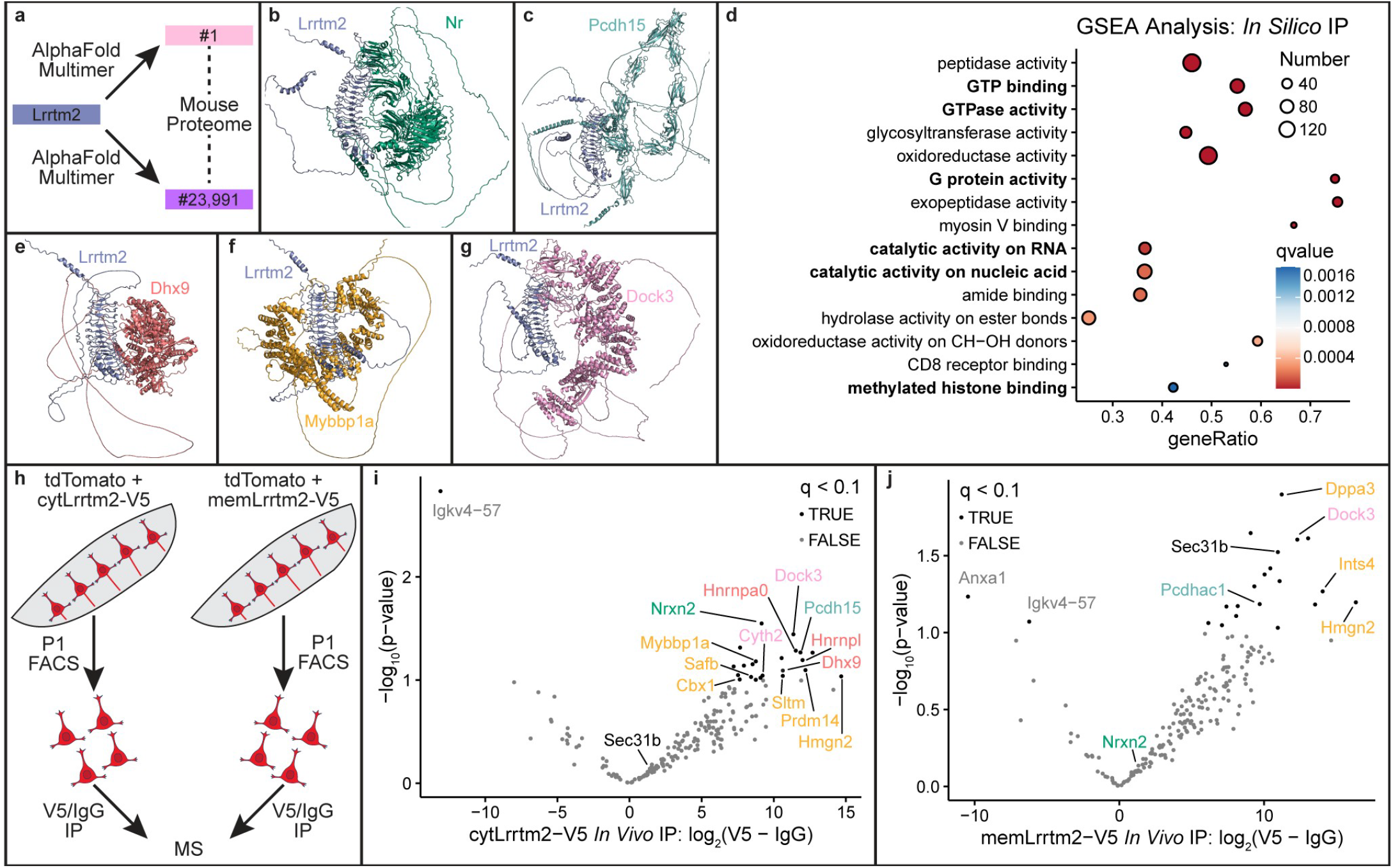
Cytoplasmic Lrrtm2 binds Nrxn and other key GC proteins in CPN. **(a)** Identification of Lrrtm2 complexes via an *in silico* IP against the mouse proteome. **(b, c)** Lrrtm2 is predicted to interact with GC adhesion proteins. **(d)** GSEA of high-confidence Lrrtm2 interactors. **(e-g)** Lrrtm2 is predicted to interact with RBPs, TFs, and G proteins. **(h)** Elucidation of Lrrtm2 complexes via *in vivo* IP-MS of CPN somata at P1 after transfection with low-level cytLrrtm2 or memLrrtm2 OE constructs. **(I, j)** Proteins with significant enrichment after V5 IP relative to IgG IP for CPN with low-level Lrrtm2 OE (q < 0.1). Nrxn is green, GC adhesion proteins are teal, RBPs are red, TFs are yellow, G proteins are pink, and Sec31b is black.

To identify additional protein classes that likely interact with Lrrtm2, we performed gene set enrichment analysis (GSEA) on high-confidence Lrrtm2 complexes (Fig. 5d). Intriguingly, Lrrtm2 is predicted to bind RNA binding proteins (RBPs), TFs, and G proteins, which are known to regulate circuit formation – notable examples include RBPs Dhx9 and Cmtr2, TFs Mybbp1a and Mau2, and G proteins Dock3 and Rab11b^103–108^ (Fig. 5e-g; Supp. Fig. 7f, g, j-l, o-q). These findings indicate that cytLrrtm2 might bind many distinct proteins in CPN, and that this relatively broad proteomic dysregulation might contribute importantly to aberrant BLA innervation by cytLrrtm2^OE^ CPN *in vivo*.

To directly investigate, and potentially validate, *in silico* Lrrtm2 interactors, and to investigate differences in ligand binding between cytLrrtm2 and memLrrtm2, we performed IP-MS in CPN somata. Briefly, we transfected CPN with low-level OE constructs for cytLrrtm2 or memLrrtm2, each of which is tagged with a V5 tag, via E14.5 IUE. We purified CPN somata at P1 by FACS, and analyzed extracted proteins via IP-MS with antibodies against either V5 or a control IgG (Fig. 5h). Importantly, proteomes from these samples form distinct clusters by condition (Supp. Fig. 7r; Supp. Table 5).

Strikingly, cytLrrtm2, but not memLrrtm2, binds Nrxn *in vivo*, supporting the possibility that cytLrrtm2 sequesters Nrxn from CPN GC surfaces via direct binding (Fig. 5i, j). Although cytLrrtm2 and memLrrtm2 interact equivalently with adhesion and G proteins, cytLrrtm2 forms complexes with more RBPs (3 vs 0) and TFs (6 vs 3) than memLrrtm2. Further, memLrrtm2, but not cytLrrtm2, interacts with Sec31b, which localizes proteins with signal peptides to membranes^109^. Together, these results reveal that aberrant localization of Lrrtm2 to the cytoplasm of CPN, rather than to the membrane, leads to binding of many molecules that control precise circuit formation, including sequestration of Nrxn from GC surfaces, causing aberrant BLA innervation.

## Discussion

### cytLrrtm2 sequesters GC proteins to dysregulate wiring

Here, we discover that non-canonical Lrrtm2 sequesters GC surface proteins to dysregulate CPN circuit wiring. As a first step, we identify that Lrrtm2, a canonically postsynaptic transmembrane protein, also functions in developing GCs to regulate circuit formation and specificity. We identify this previously unknown control over precise circuit wiring by elucidating the function of dysregulated Lrrtm2 in causing *Bcl11a*^-/-^ CPN to aberrantly target and innervate the BLA. Importantly, proteomes from these samples form distinct clusters by condition (Supp. Fig. 7r; Supp. Table 5).

We investigate and identify dysregulated subcellular proteomes of *Bcl11a*^-/-^ CPN via optimized approaches for *in vivo* subtype-specific GC isolation and ultra-low-input mass spectrometry (Fig. 1; Supp. Fig. 1; Materials and Methods). We verify localization of several candidate proteins to CPN GCs *in vivo* by further developing subcellular approaches (Fig. 2; Supp. Fig. 2), and we deeply investigate functions of an exemplar: Lrrtm2. These results reveal that cytLrrtm2, but not memLrrtm2, cell-autonomously causes aberrant innervation of BLA by *Bcl11a*^-/-^ CPN (Fig. 3; Supp. Fig. 3).

We further identify that cytLrrtm2, but not memLrrtm2, sequesters Nrxn from CPN GC surfaces *in vivo* via recently engineered approaches (Fig. 4; Supp. Fig. 7). Finally, we identify that cytLrrtm2, but not memLrrtm2, directly binds Nrxn and other key GC proteins, and find that cytLrrtm2 interacts with other proteins in CPN, via *in vivo* and *in silico* IP approaches (Fig. 5; Supp. Fig. 7), likely inhibiting their normal functions. Together, these results identify a non-canonical molecular mechanism of cytoplasmic GC protein sequestration regulating *in vivo* circuit development, reveal how control over localized subcellular GC proteomes is required for precise subtype-specific circuit formation, and highlight how dysregulation of these intricate processes drives aberrant wiring, dysfunction, and human disorders.

### Non-canonical control over GC surface proteomes *in vivo*

Prior work has identified GC receptors that regulate axon guidance by integrating information from extracellular cues to control cytoskeletal remodeling, and have demonstrated that insertion of these proteins into membranes is required for their function^110,111^. To our knowledge, however, there are no examples of key GC surface proteins being bound and sequestered by a cytoplasmic GC protein for control over membrane insertion and function, and thus precise circuit formation. Considering the importance of GC receptors in regulating axon growth and guidance, combined with the results presented here, it is quite likely that binding and sequestration of GC surface proteins by cytLrrtm2 and/or other cytoplasmic molecules contributes to construction of typical and aberrant circuitry across neuron subtypes. Our recently developed experimental and analytical approaches enable further *in vivo* investigation of such non-canonical molecular mechanisms in distinct circuits at distinct stages, and potential identification of how this subtle dysfunction disrupts functional circuitry. We speculate that these non-canonical mechanisms might contribute to powerful and generalizable regulation over timing, stages, and transitions of circuit development, maintenance, and (dys)function.

In addition to regulating GCs directly, mislocalization of Lrrtm2 to cytoplasm might impact circuit wiring via binding of distinct non-surface proteins. Our *in silico* and *in vivo* IP experiments indicate that cytLrrmtm2 selectively interacts with other controls over neural development, including TFs (e.g. Cbx1^112^), RBPs (e.g. Hnrnpa0^113^), the adhesion protein Pcdh15^114^, and the G protein Cyth2^115^. However, it remains unknown how function of these proteins might be regulated by cytLrrtm2 binding and potential sequestration, or how such disruptions might affect circuit development, function, and maintenance. Furthermore, localization of Lrrtm2 to cytoplasm likely prevents its canonical insertion into CPN postsynaptic membranes. Since Lrrtm2 has roles in synapse formation and modulation^54,55,79,116^, absence of this protein from dendritic membranes downstream of *Bcl11a* might disrupt activity of CPN, including local CPN subnetworks and homotopic connections with contralateral CPN^117,118^.

### A molecular regulator of BLA targeting and innervation

Despite the importance of BLA in regulating affect, social behavior, cognition, and anxiety, its hyperactivation in ASD, and its dysregulated connectivity with cerebral cortex in bipolar disorder^38–40,119,120^, little is known about molecular controls regulating BLA innervation. For example, recent work has identified that Efnb3 controls BLA targeting by CA1 hippocampal neurons^121^, but it is unclear how other BLA-projecting populations (e.g. piriform cortex, prefrontal cortex, thalamus^122,123^, as well as *Mef2c*^-/-^ CPN^25^) specifically target BLA. Since cytLrrtm2^OE^ CPN project to BLA, and since other leucine-rich repeat proteins regulate construction of distinct circuits^81–84^, it is possible that localization of Lrrtm2 to cytoplasm is a general mechanism for innervation of this ancient brain region by a range of diverse neuron subtypes.

Although we identify that Lrrtm2 is more abundant in *Bcl11a*^-/-^ CPN GCs that are extending through contralateral cortex at P3, and demonstrate that cytLrrtm2^OE^ CPN target BLA, many *Bcl11a*^-/-^ and cytLrrtm2^OE^ CPN still innervate contralateral cortex, rather than BLA. This result strongly suggests additional molecular and sub-subtype diversity within CPN that is mostly undescribed. Our lab has made progress towards delineating distinct subpopulations of CPN^124–126^, and recent work identifies subtle, but critical, differences in GC proteomes between medial and lateral CPN^16^. However, little is known about how wild-type CPN subpopulations innervate multiple targets, how some CPN extend dual projections to distinct regions^126,127^, and how *Bcl11a* deletion leads to distinct circuit disruptions within CPN. Further investigations pairing subcellular purification with ultra-low-input transcriptomics and proteomics has deep promise to resolve these questions, and potentially reveal important molecular, functional, and circuit diversity across distinct subpopulations of sparse CPN sub-subtypes.

### Subcellular approaches identify distinct circuit regulators

Despite the power of cell body transcriptomics to elucidate subtype-specific controls over circuit wiring^3^, many other regulators are “missed” via this more traditional approach. Similarly, elegant *in vitro* studies have identified molecular networks that are required for GC function broadly^10^, but they do not optimally reproduce precise subtype identities, nor the complexity of *in vivo* extracellular environments. In contrast, subcellular-and subtype-specific investigations *in vivo* are well-positioned to elucidate GC molecular networks at select stages of circuit formation, if sufficient material is available. Transcriptomic investigation requires very little input and enables identification of RNAs that are translated in GCs, but does not capture active protein function, since GCs synthesize proteins locally in response to extracellular cues, and also employ proteins that were translated in cell bodies^47–49^. Thus, subtype-specific subcellular proteomics *in vivo* is highly likely to identify distinct wiring regulators.

The enhanced approaches for purification of subcellular compartments and ultra-low-input mass spectrometry that we employ here will enable continued identification of *bona fide* GC molecules across neuron subtypes and distinct stages of development, maintenance, and (dys)function. These improvements in sensitivity and accuracy will also benefit transcriptomic approaches, and have promise to accelerate design of subcellular ribosome profiling for more direct investigation of local translation in subtype-specific GCs^62^. Similar principles can be applied across subcellular compartments (e.g. synapses, dendrites), further deepening understanding of how neurons and other polarized cell types regulate subcellular localization and local function.

### GC molecules control specific aspects of circuit formation

*BCL11A* is a fully penetrant, monogenic ASD/ID-risk gene in humans, and is associated with bipolar disorder^32–36,128,129^. Prior work from our lab and others revealed that mice with *Bcl11a*^-/-^ CPN exhibit defects in CPN circuit formation, cell body transcription, and social behaviors^29–31^. More recent work from our lab expanded understanding of how *Bcl11a* regulates soma gene expression, circuit development, and subcellular GC RNA localization, and elucidated function of downstream GC RNAs controlling precision of contralateral innervation^28^ (Fig. 6). In this work, we substantially deepen understanding of *Bcl11a* function by identifying disrupted subcellular proteomes in *Bcl11a*^-/-^ CPN, and discover a non-canonical mechanism underlying the strikingly aberrant, *de novo*, and highly specific BLA innervation by *Bcl11a*^-/-^ CPN. This work demonstrates that Lrrtm2, and likely Sema3f, function downstream of *Bcl11a* in CPN to regulate aberrant BLA targeting and midline crossing, respectively. Strikingly, these GC molecules control distinct aspects of CPN wiring (e.g. interhemispheric targeting, BLA circuitry, axon growth rate), indicating that *Bcl11a* deletion disrupts independent molecular networks, and that multiple orchestrated axon guidance programs enable precise wiring, as others have suggested^130^. Notably, manipulating each downstream GC molecule generates a specific CPN circuit defect, rather than contributing to each of the multiple disruptions of *Bcl11a*^-/-^ CPN connectivity. This separability of distinct circuit wiring “modules” enables future investigation of how these circuit elements arise for proper function, and how even subtle perturbations may disrupt specific social behaviors in mice.

**Figure 6:**
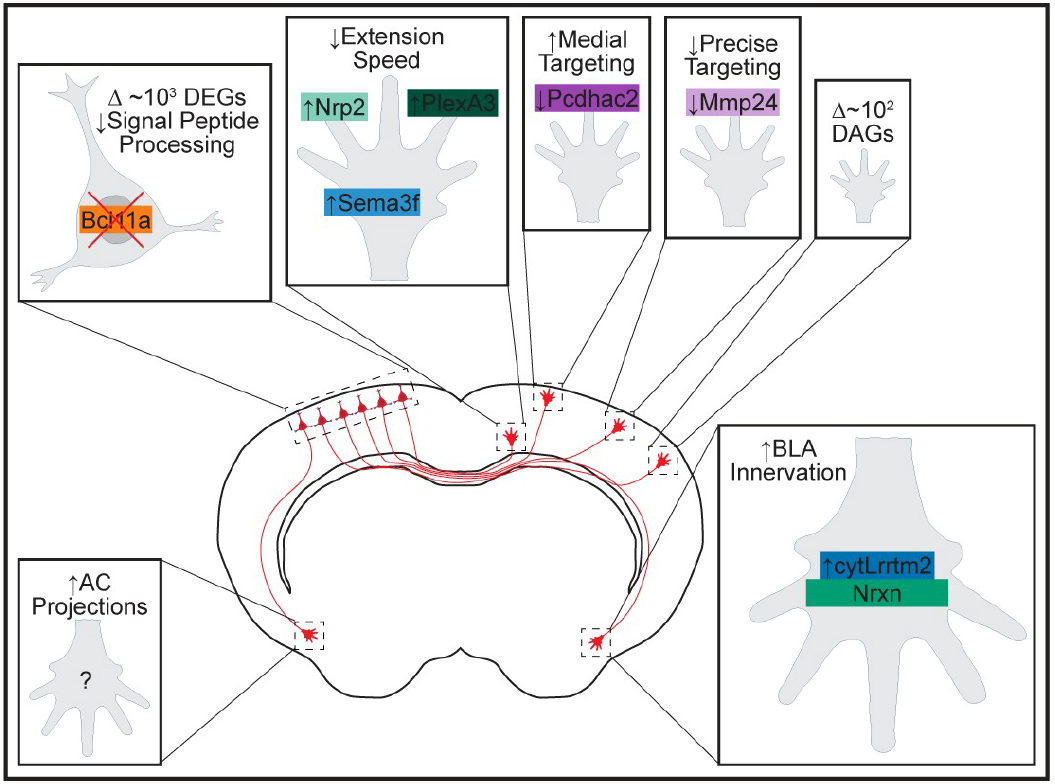
*Bcl11a* regulates diverse molecular networks to control precise CPN circuit formation. *Bcl11a*^-/-^ CPN have dysregulated gene expression and subcellular localization of RNAs and proteins. Specific downstream molecules regulate distinct aspects of CPN circuit wiring. Molecular controls over increased AC projections remain unknown.

More broadly, our finding that signal peptide processing is dysregulated by *Bcl11a* loss-of-function indicates that other membrane-inserted and/or secreted proteins might be mislocalized in *Bcl11a*^-/-^ CPN. Strikingly, all identified GC molecules downstream of *Bcl11a* normally contain signal sequences^28^, and membrane proteins are key regulators of circuit formation, maintenance, and function across neuron subtypes^10^. Therefore, it is highly likely that at least some non-CPN cell types aberrantly localize membrane proteins in humans with *BCL11A*-related ASD/ID, since *BCL11A* is variant or mutated across all cells of these patients. Beyond *BCL11A* variants, it is possible that this dysregulation of membrane proteins (e.g. membrane insertion, subcellular localization) contributes to other forms of ASD and ID, as well as other neurodevelopmental disorders across a range of key nervous system circuits. Ongoing work will further elucidate how loss of TF function dysregulates localization of molecules across distinct subcellular compartments, and their insertion into membranes, and how these disruptions cause dysgenesis of normally precise and distinct circuits *in vivo*, resulting in atypical function and behaviors.

## Acknowledgements

We thank Jaewon Heo and Melody Ross for technical support; members of the Macklis lab for insightful suggestions; Nema Kheradmand, Gracia Kassis, and Joyce LaVecchio of the HSCRB-HSCI Flow Cytometry core; and Claire Hartmann, Nicole Ramirez, and Christian Daly of the Bauer Core Facility. This work was supported by the following grants to J.D.M.: NIH DP1 NS106665, NIH R01 NS104055; additional infrastructure support and technology from the Eagles Autism Foundation, and NIH R01 NS045523, NIH R21 NS141110, NIH RF1 AG083085, NIH R21 NS104733, and Max and Anne Wien Professor of Life Sciences fund. D.E.T. was partially supported by NIH F31 NS127518, and the NSF-Simons Center for Mathematical and Statistical Analysis of Biology at Harvard #1764269. O.D. was partially supported by NIH T32 AG000222. P.V. was partially supported by NSF GRFP 280932, the NSF-Simons Center for Mathematical and Statistical Analysis of Biology at Harvard #1764269, and NIH T32 GM007306-43. J.E.F. was partially supported by NIH T32 AG000222, and NIH F32 AG067661. This paper was typeset with the bioRxiv word template by @Chrelli: www.github.com/chrelli/bioRxiv-word-template

## Author contributions

D.E.T. and J.D.M. conceived the overall project and experiments. D.E.T. designed individual experiments and performed all experiments unless otherwise noted; O.D. performed *in utero* electroporations for mass spectrometry experiments; P.V. performed *in utero* electroporations and read mapping for growth cone “washing” experiments, and assisted with sample collection for soma mass spectrometry; J.E.F. assisted with sample collection for soma mass spectrometry; G.W. performed sectioning, labeling, and imaging; B.B. performed mass spectrometry. D.E.T., and J.D.M. analyzed and interpreted data, and wrote and edited the manuscript.

## Competing interest statement

The authors declare no competing interests.

## Materials and Methods

### Mice

All mouse work was approved by the Harvard University Institutional Animal Care and Use Committee (IACUC), and performed in accordance with institutional and federal guidelines. Outbred CD1 mouse pups were ordered from Charles River Laboratories (Wilmington, MA). *Bcl11a*^fl/fl^ mice were generated and generously shared by H. Tucker and G. Ippolito^131,132^. All *Bcl11a*^fl/fl^ mice were maintained on a CD1 background. The morning of vaginal plug identification was designated as E0.5, and the day of birth as P0. Mice were housed in groups of 2-4, except males that were single housed post-breeding, all on a standard 12 h light / 12 h dark cycle.

### *In utero* electroporation

Unilateral IUE was performed at E14.5, as we and others have previously described^133^. Briefly, pregnant females were anesthetized using isoflurane (VetOne #502017), then lower abdominal hair was removed, and skin was locally disinfected. An incision was made to expose the uterine horns, which were gently externalized from the intraperitoneal cavity onto a moist, sterile gauze pad. Lateral ventricles were unilaterally injected with plasmid mixtures containing Fastgreen (Thermo #AAA16520-06) using a beveled glass micropipette (Fisher #22-260-943). Five 34 volt pulses (50 ms on and 950 ms off per pulse) were delivered with 5 mm diameter platinum plate electrodes (Protech CUY650-P5), and a CUY21EDIT square wave electroporator (Nepa Gene). Embryos were then gently placed back into the intraperitoneal cavity, and the wound was closed with sutures (Covetrus #39010). The pregnant dam fully recovered from anesthesia and surgery on a thermally-controlled mat under low-intensity heat before being returned to animal husbandry. Postoperative care included daily checks of well-being and the wound, and administration of buprenorphine (Par Pharmaceutical #60969). After birth, injected pups were screened for unilateral fluorescence using a fluorescence stereoscope. Concentrations of plasmids across experiments are listed below. Fluorophore-only: 3 ug/ul of fluorophore plasmid; Bcl11a KO: 3 ug/ul of fluorophore plasmid and 1 ug/ul of Cre plasmid; OE: 3 ug/ul of fluorophore plasmid and 1 ug/ul of OE plasmid; BEAM: 2 ug/ul Cre-On fluorophore with OE protein plasmid, 2 ug/ul Cre-Off fluorophore plasmid, 0.5 ug/ul Cre-On Cre plasmid, 10-50 ng/ul Cre plasmid (optimized for each maxiprep to obtain 50:50 red/green ratios via FACS). All constructs and oligos are listed (Supp. Tables 6, 7).

### Soma purification via fluorescence-activated cell sorting

Fluorescent cell bodies were purified as we have previously described^11,12^. Briefly, brains were extracted into cold HBSS (Thermo #14025134), and layer II/III CPN were microdissected into dissociation solution (DS). Tissue chunks were washed twice with DS, then enzymatically dissociated via two 15 minute incubations with enzyme solution supplemented with DNaseI (Worthington #LS006330). Dissociated tissue was washed twice with wash solution (WS), triturated with a fire-polished Pasteur pipet, washed with WS, and pelleted at 80*g* for 5 minutes at 4 °C. Cells were resuspended in WS, and CPN were purified via FACS, using a custom BD SORP Aria II. qPCR was performed via SuperScript IV reverse transcriptase (Thermo #18090050) and SsoAdvanced Universal SYBR Green Supermix (BioRad #1725271), using a QuantStudio3 machine (Thermo). Code is provided: github.com/detillman/lrrtm2_macklis_2026.

### Growth cone purification via fluorescent small particle sorting

Fluorescent CPN GCs were purified by optimizing approaches that we have previously described^11,12^. Briefly, ipsilateral cortical hemispheres were microdissected into cold 0.32 M sucrose containing 4 mM HEPES, (Thermo #15630106), Halt protease inhibitor (Thermo #78442), and RNasin RNase inhibitor (Promega #N2515), then homogenized at 900 RPM in a glass-Teflon tissue grinder. Post-nuclear homogenate (PNH) was generated by collecting supernatant after centrifuging at 1,660*g* for 15 min at 4 °C, and PNH was layered onto a 0.83 M and 2.5 M sucrose gradient. The sucrose gradient was centrifuged at 242,000*g* for 47 min at 4 °C in a vertical rotor (Beckman VTi50), enabling isolation of bulk GC fraction (GCF) via extraction of the interface between 0.32 M and 0.83 M sucrose. Fluorescent GCs were purified from dilute GCF, supplemented with 0.5 um Fluoresbrite YG beads (Polysciences #17152) for some optimization experiments, via FSPS, using a customized BD SORP Aria II (Supp. Fig. 8a-c).

### Growth cone protection assay

GC protection assays were performed as we have previously described^11,12^. For protein protection assays, 40 μl of GCF was rocked at 4 °C for 90 min with 1 μl 20% Triton (Sigma X100) and/or 8 μl 25% trypsin (Thermo #25200056) or nothing. Loading buffer (Thermo #39001) and beta-mercaptoethanol (Sigma #M6250) were then added to all samples, which were incubated at 95 °C for 5 min, and frozen at-20 °C prior to western blot. For RNA protection assays, 60 μl of GCF was rocked at 4 °C for 90 min with 1 μl 20% Triton (Sigma X100) and/or 1 μl 20% RNase ONE (Promega # M4261) or nothing. RNA was extracted from all samples (Zymo #R1050), and analyzed via TapeStation (Agilent #G2991BA). RNA is present in bead samples, and GCs co-purified with beads are protected (Supp. Fig. 8d-f).

### Western blot

Samples were loaded into wells of a 4-20% Mini-PROTEAN TGX precast protein gel (Biorad #4561096), and run at 150 V for 45 min. Gels were transferred via Trans-Blot Turbo Mini 0.2 um PVDF transfer packs (Biorad #1704156) at 2.5 A for 7 min, washed with TBS-T for 5 min, blocked with 5% milk in TBS-T for 1 hour, rinsed with TBS-T twice, washed with TBS-T for 15 min, and incubated with primary antibody in 5% BSA at 4 °C overnight. Membranes were washed three times with TBS-T for 10 min each, incubated with secondary antibody in 5% milk at RT for 1 hour, washed three times with TBS-T for 10 min each, and visualized with SuperSignal West Pico PLUS (Thermo #34580) using a CCD camera imager (FluoroChemM, Protein Simple). A table of all antibodies used, and their concentrations across experiments, is provided (Supp. Table 8).

### Identification of ambient RNAs in purified growth cone samples

CPN GC transcriptomes that were previously obtained by homogenizing a mixture of B6 brains, containing red CPN via E14.5 IUE, with unlabeled CD1 brains prior to subcellular fractionation and FSPS were analyzed with a custom pipeline using GATK HaplotypeCaller (v4.2.5.0)^134^. GO enrichment analysis was performed with ClusterProfiler^135^, and similar GO terms were reduced based on semantic similarity via rrvgo^136^. Increasing CD1 tissue mass leads to more CD1 SNPs in RNAs in purified CPN GCs, and RNAs without CD1 SNPs are enriched for GC GO terms (Supp. Fig. 8h, i; Supp. Table 9). Code is provided: github.com/detillman/lrrtm2_macklis_2026.

### Removal of ambient molecules from purified growth cone samples

Based on prior GC and synaptosome isolation protocols^137–139^, washed GCFs (wGCFs) were obtained via 1:10 dilution of GCF with 0.32 M sucrose, centrifugation at 15,000*g* for 30 minutes at 4 °C, removal of supernatant, and resuspension of pelleted GCs in 1 mL of 0.32 M sucrose (Supp. Fig. 8j). wGCFs retain Gap43, are depleted of Map2, and are protected (Supp. Fig. 8k-n). Washed GCs (wGCs) were purified from wGCFs by homogenizing a mixture of CD1 brains, containing red CPN via E14.5 IUE, and unlabeled B6 brains prior to subcellular fractionation and FSPS (Supp. Fig. 9a). RNA is absent from beads co-purified from wGCFs (Supp. Fig. 9b). RNA was extracted from wGCs, GCs, and supernatants (supers) with the Allprep Mini Kit (Qiagen #80204), and libraries were prepared via SmartSeq v4 (Takara Bio) ¼-volume reactions, then a NovaSeq (Ilumina) was used for 2×100 bp paired-end reads for full-transcript sequencing of ∼25-40M reads per sample. Adapters were clipped and unpaired reads were trimmed with Trimmomatic (v0.39). Reads were aligned to the GRCm39 mouse genome with STAR (v2.70e) in 2-pass mode^140^, exonal reads were quantified with featureCounts (v1.5.1)^141^, and SNPs were identified with a custom pipeline (see above). wGCs have ∼5-fold more high-confidence CD1 SNPs than GCs or supers, and form a distinct PC cluster (Supp. Fig. 9c, d; Supp. Table 10). Differentially abundant RNAs were identified via DESeq2^142^, effect sizes were shrunk via apeglm^143^, and counts were transformed via rlog^142^. wGCs are depleted of RNAs relative to GCs and supers – these RNAs contain CD1 SNPs, and are enriched for GC GO terms (Supp. Fig. 9e-h; Supp. Table 11). wGCFs and wGCs were used in all other experiments (Fig. 1-5; Supp. Fig. 1-7). Counts are on Harvard Dataverse (doi.org/10.7910/DVN/KZ2JDL), and code is provided: github.com/detillman/lrrtm2_macklis_2026.

### Mass spectrometry of somata and growth cones

CPN GCs and somata were sorted via FSPS or FACS into protein LoBind tubes (Eppendorf #022431081), after tubes were rinsed with acetonitrile (Sigma #34851), containing nuclease-free water (Thermo #4387936) or 50 mM TEAB pH 8.5 (Sigma #90360), respectively, concentrated to 100 μl via centrifugation in 10 kDa concentrators (Millipore #UFC8010), and frozen at-80 °C. Samples were then thawed, digested by Trypsin Platinum (Promega #VA9000), and run on an Orbitrap Astral mass spectrometer (Thermo #BRE725600). GC samples, which contain sucrose, were diluted with TEAB prior to “one-pot” digestion, injected by a Vanquish Neo UHPLC (Thermo #VN-S10-A-01) onto a Double nanoViper PepMap trap column (Thermo #DNV75150PN), extensively washed with 0.1% formic acid, and eluted via a PepSep C18 analytical column (Bruker #1893625). Soma proteins were identified via ProteomeDiscoverer and Chymerys, and GC proteins were identified via Comet^144^ after converting.raw files with ProteoWizard^145^. GC and soma proteomes were both analyzed with DEP^146^. Proteomes are on Harvard Dataverse (doi.org/10.7910/DVN/KZ2JDL), and code is provided: github.com/detillman/lrrtm2_macklis_2026.

### Immunofluorescence of adherent growth cones

12 mm glass coverslips (Fischer #12-541-001) were incubated with a solution of 1% gelatin (Sigma #G9382) and 1% chromium potassium sulfate (Sigma #243361) in a 24-well plate for four minutes, then dried at 60 °C for 2 hours. 100 μl of wGCF was diluted with 900 μl of 0.32 M sucrose, and added to each coverslip for adhering via centrifugation at 2,250*g* for 82 minutes at 4 °C. Supernatant was removed, and GCs were fixed in 400 μl 4% PFA and 4% sucrose for 10 minutes at RT. GCs were washed 3 times with 500 μl PBS for 5 minutes each, and stored in PBS at 4 °C overnight. Coverslips were blocked and permeabilized with 500 μl PGT (2 g/L gelatin, 0.25% Triton in PBS) for 1 hour, incubated with primary antibodies for 1 hour, washed three times with PGT for 5 minutes each, incubated with secondary antibodies for 1 hour, washed three times with PGT for 5 minutes each, washed with PBS for 5 minutes, washed with water for 5 minutes, and mounted onto glass slides. Images were acquired via an epifluorescence microscope with a motorized stage and 40x oil objective (Nikon NiE), and stitched via NIS Elements (Nikon). GC colocalization was quantified via custom ImageJ scripts, adapted from prior work with synaptosomes^77^, and FlowJo. A table of all antibodies used, and their concentrations across experiments, is provided (Supp. Table 8), and code is provided: github.com/detillman/lrrtm2_macklis_2026.

### Immunocytochemistry

Anesthetized mice were transcardially perfused with PBS and 4% PFA, and extracted brains were incubated in 4% PFA at 4 °C overnight. Brains were then washed with PBS, and cryoprotected in 30% sucrose at 4 °C prior to embedding in O.C.T. (Sakura #25608-930), cryosectioning (Leica, 50 um coronal sections), and storage in PBS with 0.05% sodium azide at 4 °C. Sections were then blocked with blocking solution (10% goat serum, 0.5% Triton) for 1 hour, and incubated with primary antibody at 4 °C overnight. Sections were incubated with secondary antibodies for 2 hours, mounted onto Superfrost glass slides (VWR #48311-702) with DAPI fluoromount (VWR #0100-01), and sealed with nail polish. Images were acquired via an epifluorescence microscope with a motorized stage and 10x objective (Nikon NiE), and stitched via NIS Elements (Nikon). BLA innervation, IUE sites, contralateral targeting, axon extension, and laminar positioning were each quantified via custom ImageJ scripts. A table of all antibodies used, and their concentrations across experiments, is provided (Supp. Table 8), and code is provided: github.com/detillman/lrrtm2_macklis_2026.

### Cell culture

HEK cells were maintained at 37 °C with 5% CO_2_ in Dulbecco’s Modified Eagle Medium (Thermo #10566024) supplemented with 10% fetal bovine serum (VWR # 97068-104) and 1% penicillin–streptomycin. Transfection was performed with Lipofectamine 3000 (Thermo #L3000001). Two days after transfection, cells were fixed in 4% PFA for 20 minutes at RT, washed twice with 0.1% BSA in PBS, and stored at 4 °C. Fixed cells were blocked with 10% goat serum and 0.5% Triton for 1 hour at RT, incubated with primary antibody at 4 °C overnight, incubated with secondary antibody at RT for 1 hour, mounted onto glass slides with DAPI fluoromount, and sealed with nail polish. Images were then acquired via a confocal microscope with a motorized stage and 60x objective (Zeiss LSM700). All constructs and oligos are listed (Supp. Tables 6, 7) and all antibodies used, and their concentrations across experiments, is provided (Supp. Table 8).

### Surface labeling fluorescent small particle sorting

slFSPS was performed as previously described^14^. Briefly, GCF was diluted 5-fold in PBS with 3% BSA, incubated with primary antibody for 1 hour at 4 °C, and incubated with secondary antibody for 20 minutes at 4 °C. GCF was then diluted 20-fold with 0.32 M sucrose, and centrifuged at 15,000*g* for 30 minutes at 4 °C. Supernatant was removed, GCs were resuspended in 1 mL 0.32 M sucrose, and analyzed via FSPS. All antibodies used, and their concentrations across experiments, is provided (Supp. Table 8), and code is provided: github.com/detillman/lrrtm2_macklis_2026.

### *In silico* immunoprecipitation

AlphaPulldown^96^ was used to predict favorable complexes with Lrrtm2 against each protein in the GRCm39 mouse proteome, using the AlphaFold database of predicted protein structures^97^. All predicted Lrrtm2 complexes with a predicted aligned error < 10 Å were analyzed via GSEA using ClusterProfiler^135^, and structures were visualized with PyMOL. Code is provided: github.com/detillman/lrrtm2_macklis_2026.

### *In vivo* immunoprecipitation

CPN somata were sorted via FACS into tubes containing N-PER (Thermo # 87792) and HALT protease inhibitor, and protein was extracted after centrifugation at 10,000*g* for 10 minutes at 4 °C. Samples were added to Protein G Dynabeads (Thermo #10004D) that had been incubated with primary antibodies for 30 minutes at 4 °C, and incubated at 4 °C overnight. Beads were washed three times with PBS-T, transferred to a new tube, and eluted in 200 mM glycine (pH 2.5), 5X SDS-PAGE loading buffer, BME, and water at 70 °C for 10 minutes. Samples were then stored at-80 °C prior to tryptic digestion, purification via S-Trap (Protifi), and analysis on an Oribtrap Astral. Proteins were then identified via Comet^144^ after converting.raw files with ProteoWizard^145^, and proteomes were analyzed with DEP^146^. All antibodies used, and their concentrations, is provided (Supp. Table 8). Proteomes are on Harvard Dataverse (doi.org/10.7910/DVN/KZ2JDL), and code is provided: github.com/detillman/lrrtm2_macklis_2026.

---

**Supplementary Figure 1:**
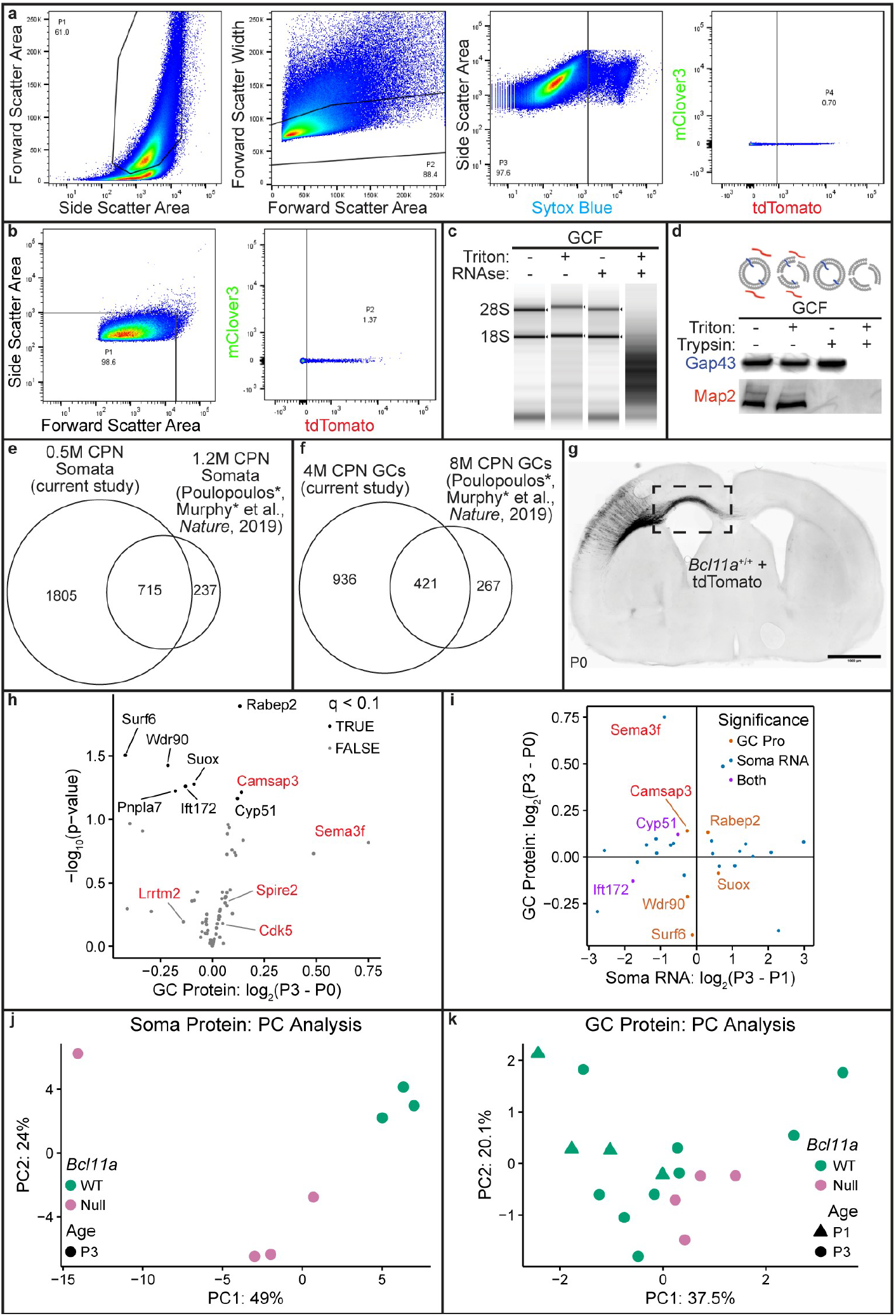
Subcellular CPN proteomics *in vivo*. **(a, b)** Representative gating strategies for FACS **(a)** and FSPS **(b). (c, d)** RNA **(c)** and protein **(d)** protection assays for GCFs. **(e, f)** Venn diagrams of protein identifications for somata **(e)** and pilot GC proteomes **(f). (h)** Proteins with differential abundance in P0 or P3 CPN GCs. Candidate protein names are red, and GC protein names are black. **(i)** Genes with differential RNA abundance in P0 or P3 CPN somata, or differential protein abundance in P0 or P3 CPN GCs. Candidate protein names are red. **(j, k)** PC analysis of soma **(j)** and GC **(k)** proteomes. Prior CPN subcellular proteomes from^11^. Soma transcriptomes from^28^. q < 0.1 for all statistical tests.

**Supplemental Figure 2:**
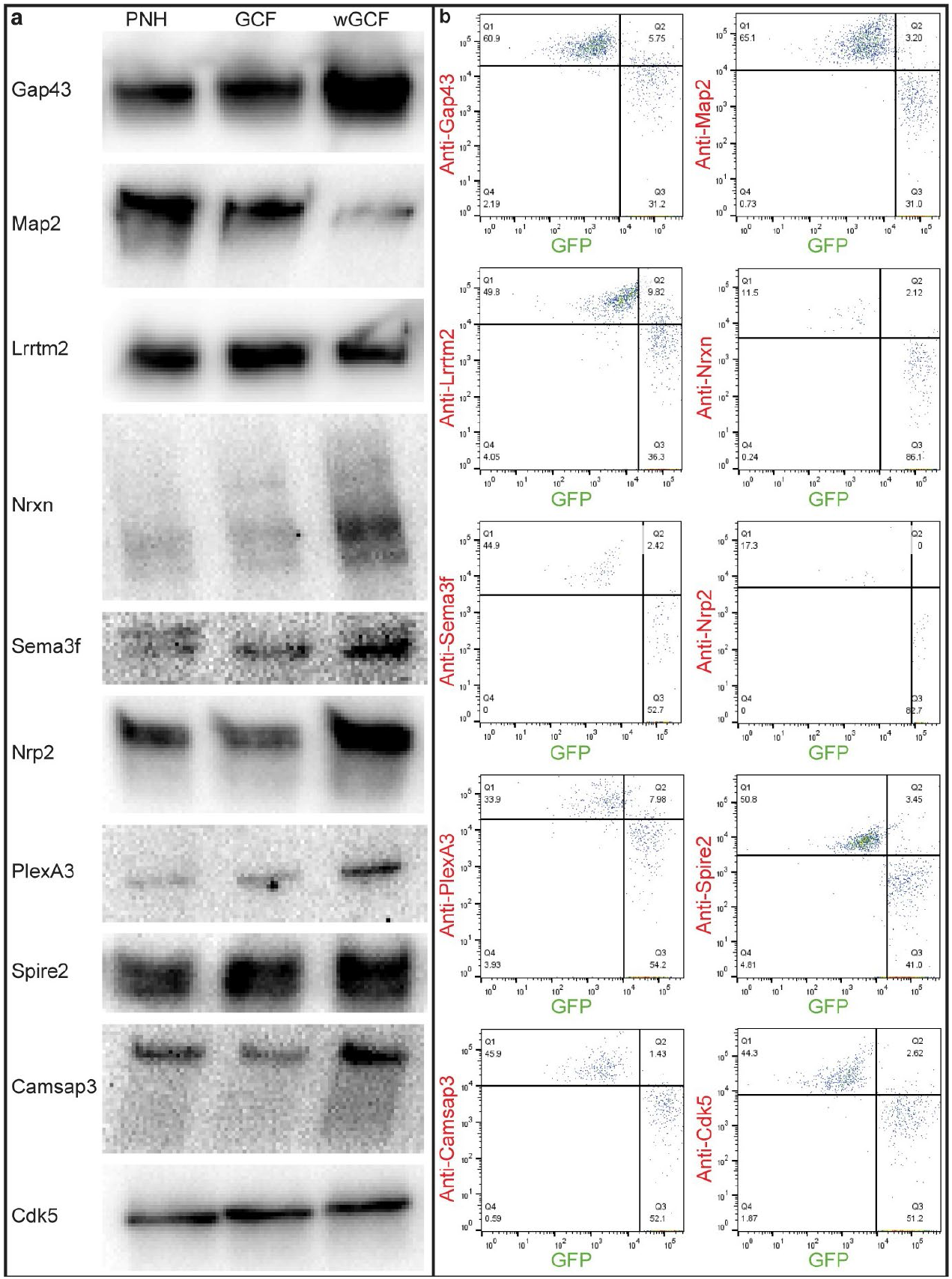
Validation of candidate GC proteins. **(a, b)** Representative western blots **(a)** and gating strategies **(b)** for GC candidates.

**Supplemental Figure 3:**
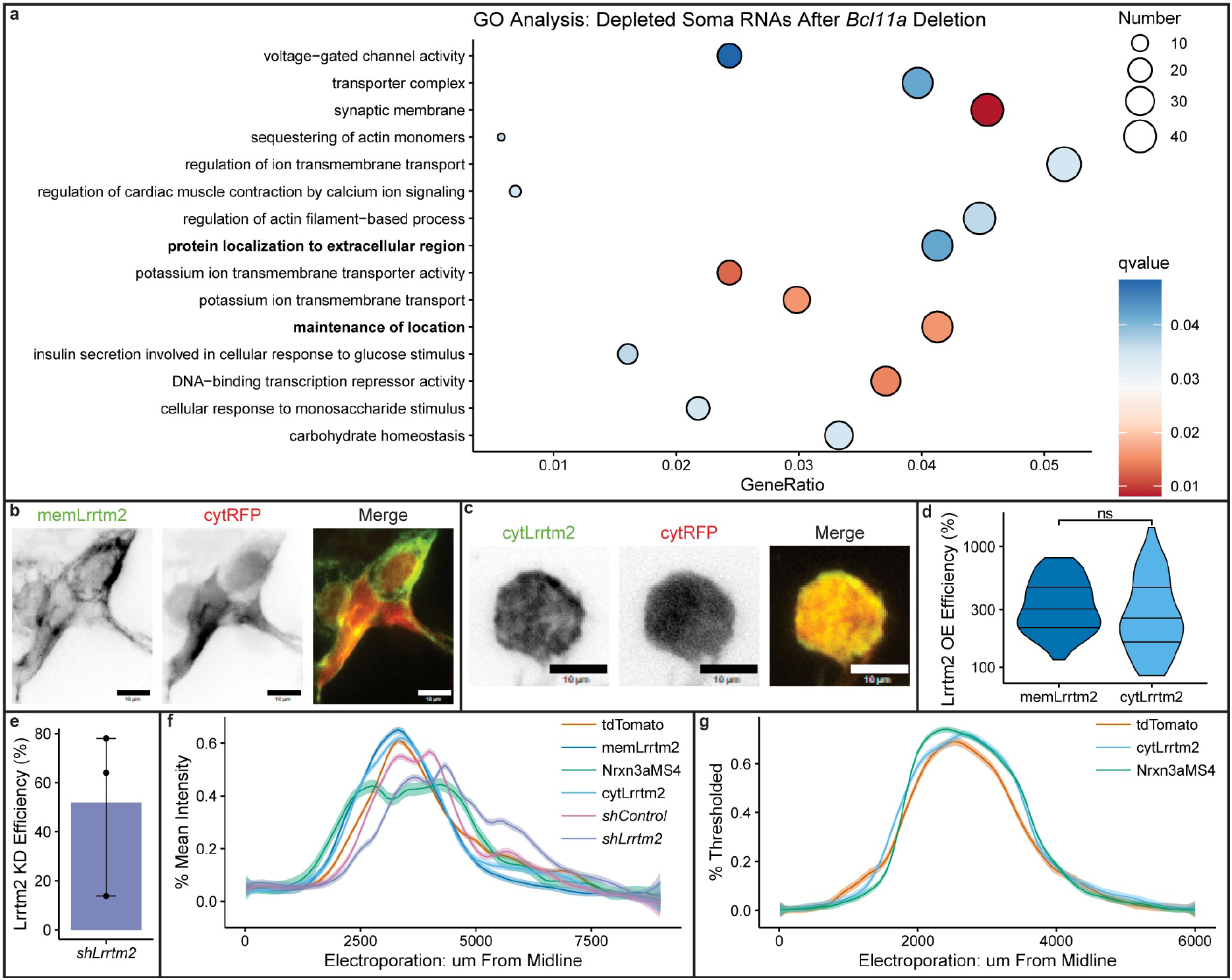
Signal peptide processing of *Bcl11a*^**-/-**^ CPN. **(a)** GO analysis of RNAs depleted in *Bcl11a*^-/-^ CPN somata. **(b, c)** HEK cells transfected with cytRFP and memLrrtm2 **(b)** or cytLrrtm2 **(c). (d)** Quantification of Lrrtm2 OE in HEK cells by microscopy. **(e)** Quantification of Lrrtm2 KD in P1 CPN somata by qPCR after E14.5 IUE of *shLrrtm2* or *shControl*. **(f, g)** IUE mediolateral locations for BLA innervation **(f)** and contralateral targeting **(g)** experiments. CPN soma transcripts from^28^. Scale bars: 10 um. Statistics: two-sided Student’s t-test (ns: p > 0.05). Error bars: 95% CI.

**Supplemental Figure 4:**
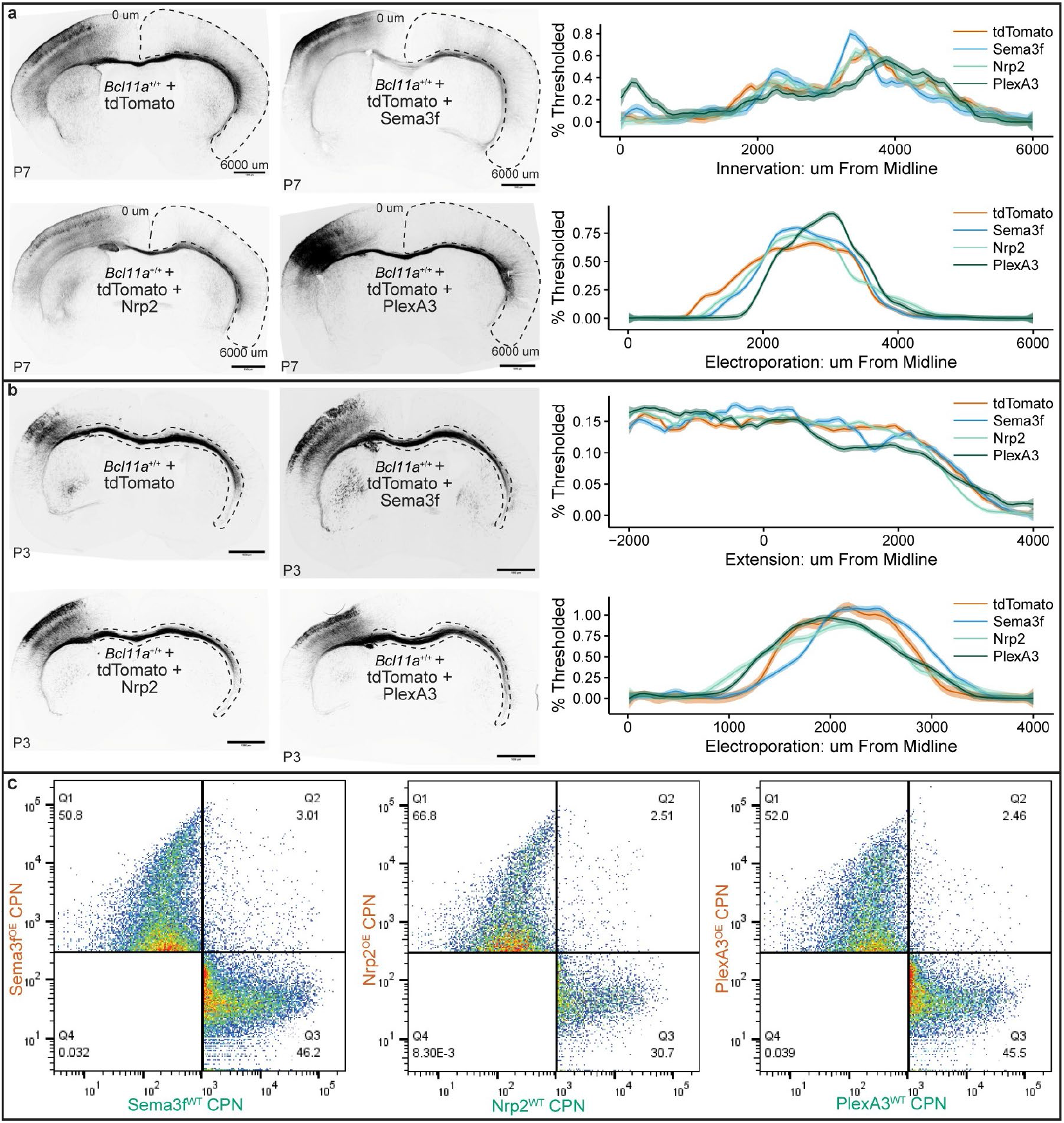
Sema3f molecular networks subtly regulate CPN circuit formation. **(a, b)** CPN are transfected via E14.5 IUE with plasmids encoding for low-level overexpression of candidate proteins, and imaged at P7 or P3 for quantification of contralateral targeting **(a)**, or axon extension **(b)**, respectively. **(c)** Representative FACS gating strategies for CPN transfected with BEAM plasmids via E14.5 IUE. Scale bars: 1000 um.

**Supplemental Figure 5:**
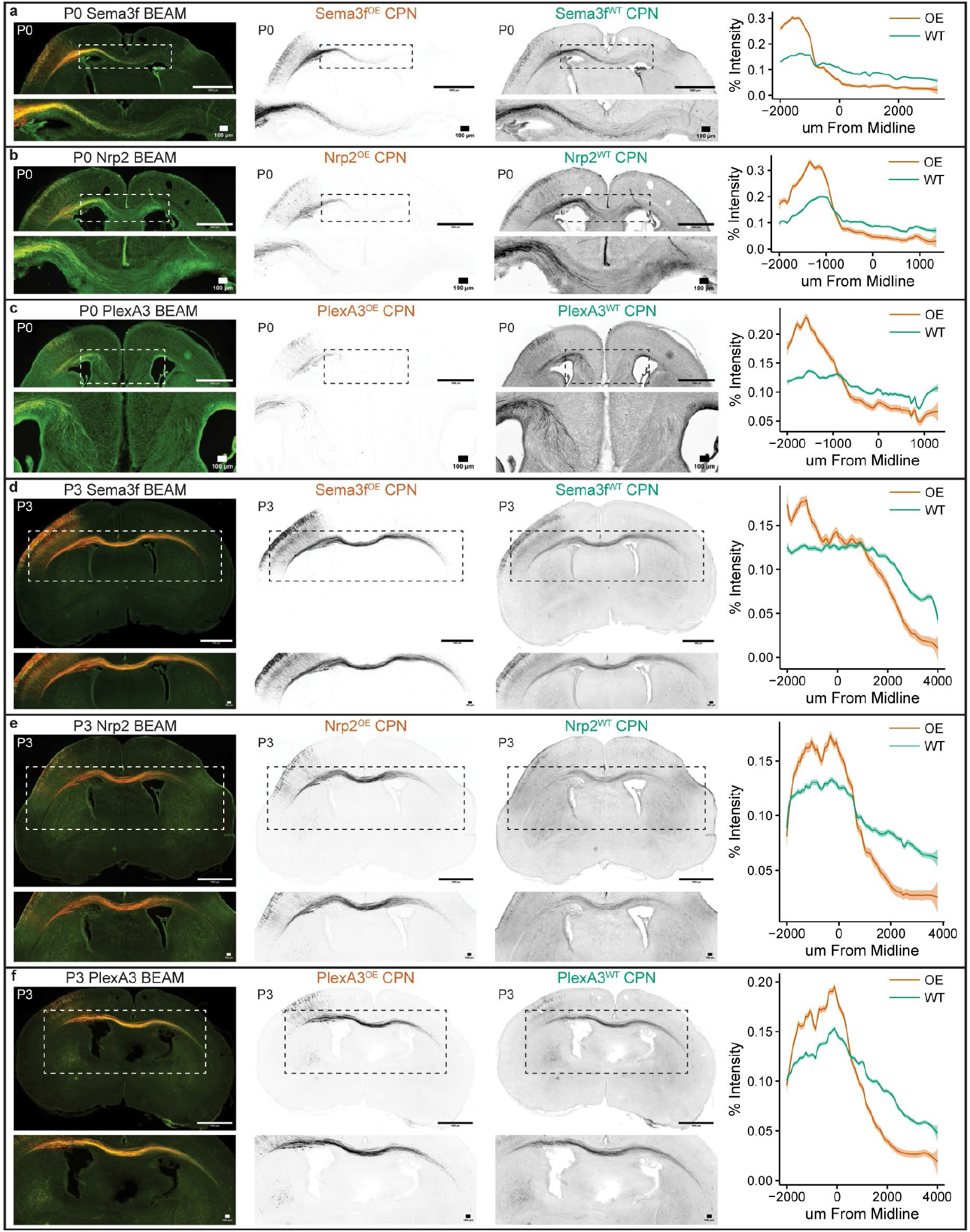
Sema3f molecular networks regulate CPN extension. **(a-f)** CPN are transfected via E14.5 IUE with plasmids encoding for BEAM Sema3f^OE^ **(a, d)**, Nrp2^OE^ **(b, e)**, or PlexA3^OE^ **(c, f)**, and imaged at P0 **(a-c)** or P3 **(d-f)**. Scale bars: 1000 um. Inset scale bars: 100 um.

**Supplemental Figure 6:**
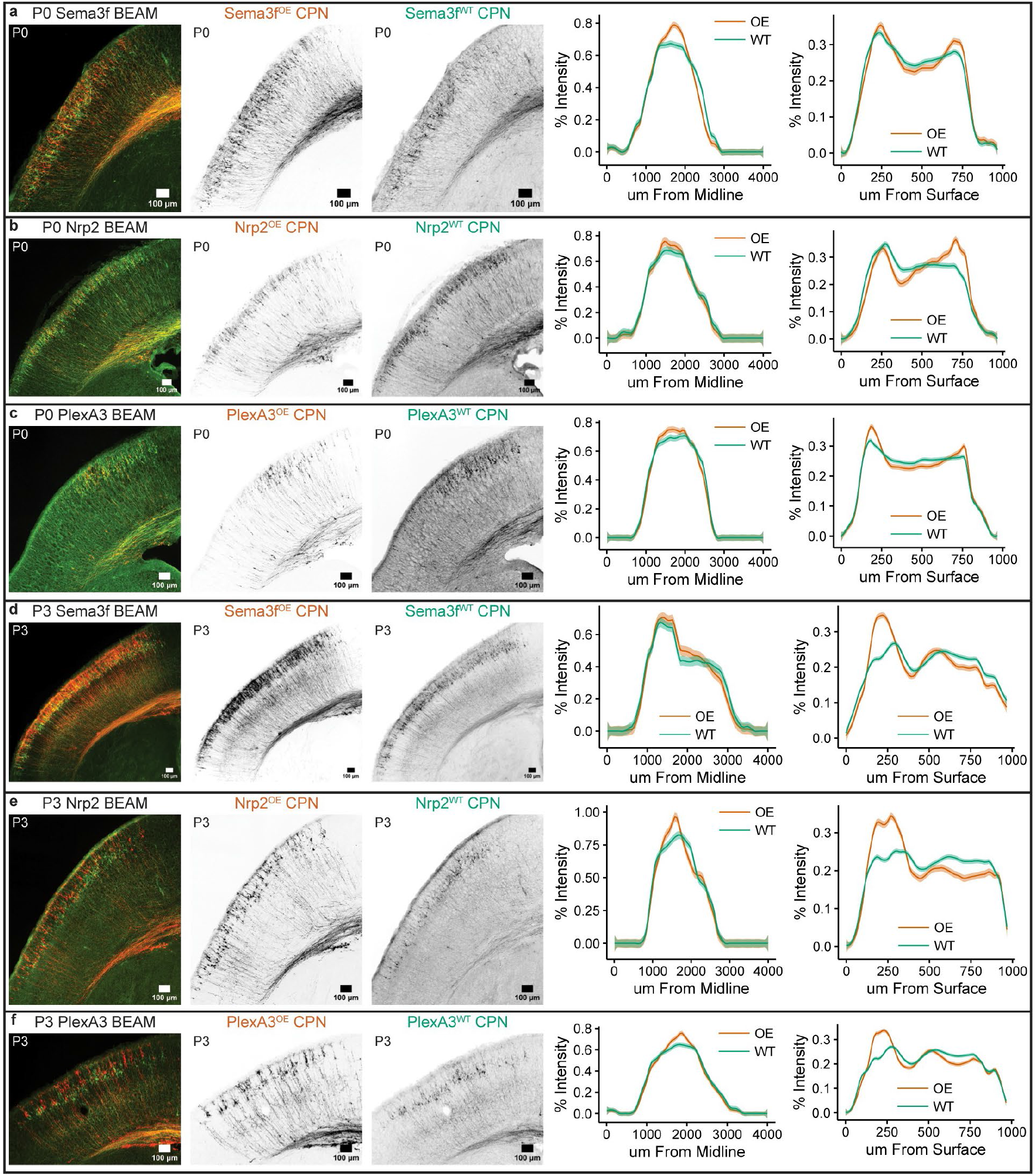
Sema3f molecular networks regulate CPN migration. **(a-f)** CPN are transfected via E14.5 IUE with plasmids encoding for BEAM Sema3f^OE^ **(a, d)**, Nrp2^OE^ **(b, e)**, or PlexA3^OE^ **(c, f)**, and imaged at P0 **(a-c)** or P3 **(d-f)**. Scale bars: 100 um. IUEs are also for Supp. Fig. 5.

**Supplemental Figure 7:**
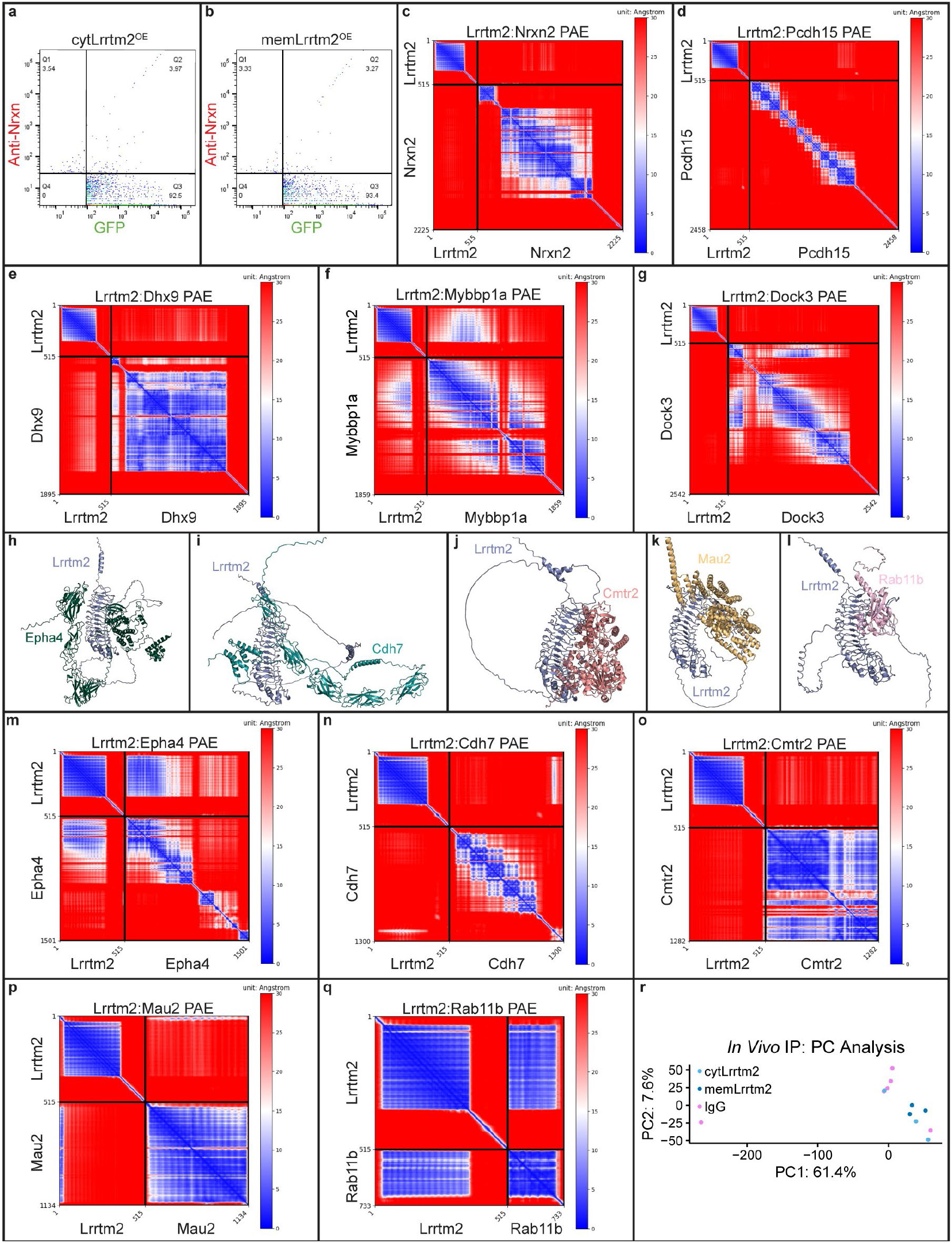
Identification of Lrrtm2 binding partners. **(a, b)** Representative slFSPS Nrxn plots for CPN transfected with low-level cytLrrtm2^OE^ **(a)** or memLrrtm2^OE^ **(b)** constructs via E14.5 IUE. **(c-g)** PAE plots for *in silico* Lrrtm2 complexes that were also identified by *in vivo* Lrrtm2 IP of CPN somata. **(h-l)** Lrrtm2 is predicted to interact with GC proteins, adhesion proteins, RBPs, TFs, and G proteins. **(m-q)** PAE plots for *in silico* Lrrtm2 complexes that were not also identified by *in vivo* Lrrtm2 IP of CPN somata. **(r)** PC analysis of *in vivo* Lrrtm2 IP proteomes from CPN somata.

**Supplemental Figure 8:**
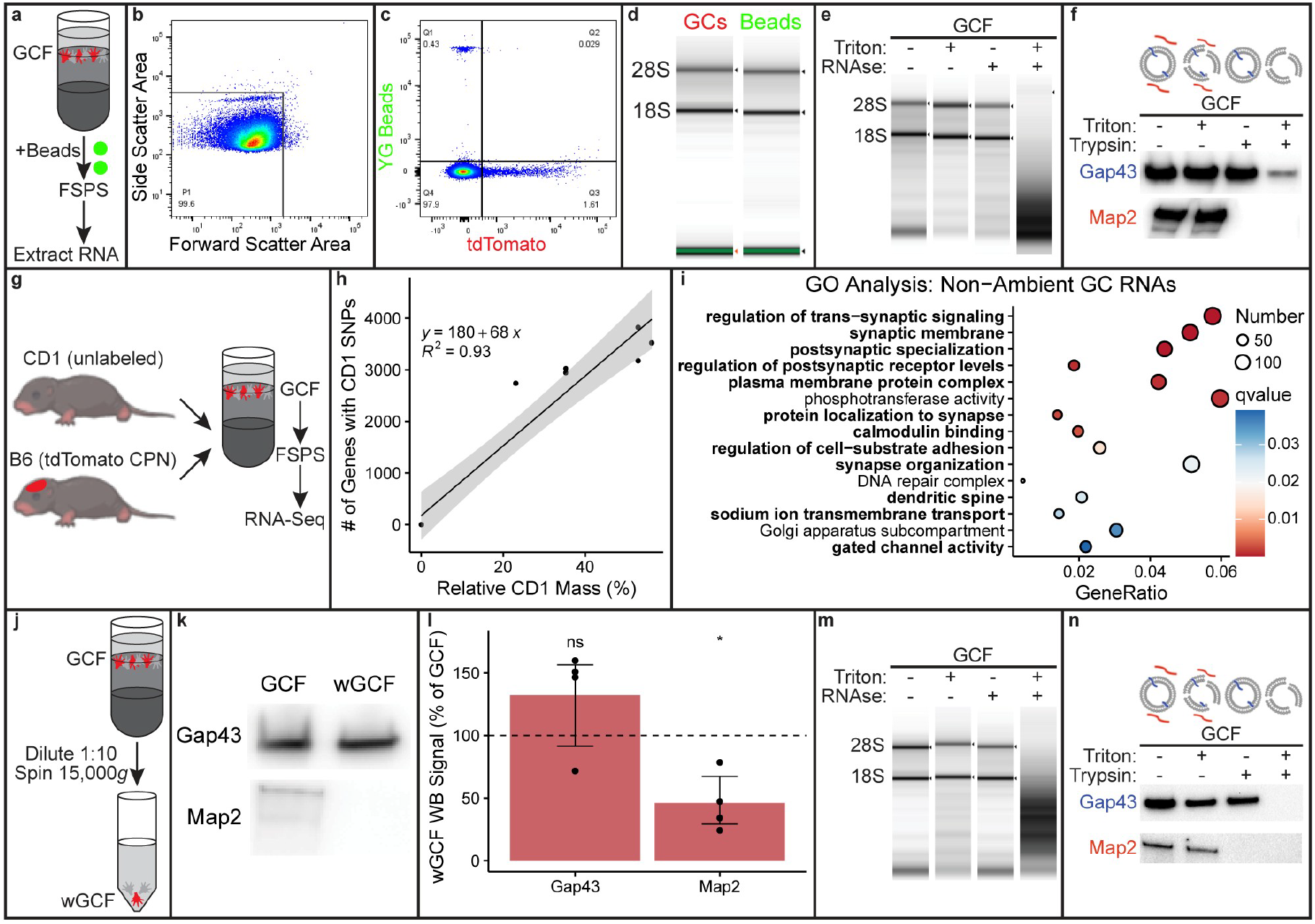
Identification of ambient molecules in purified CPN GCs. **(a)** Purifying fluorescent beads via FSPS after incubation with GCFs enables identification of environmental RNAs in GC samples. **(b, c)** Representative gating strategies for FSPS with beads. **(d)** RNA extracted from GCs and beads after FSPS. **(e, f)** RNA **(e)** and protein **(f)** protection assays for GCFs after incubating with beads. **(g)** Homogenizing unlabeled CD1 brains with B6 brains containing labeled CPN via E14.5 creates a mixed-strain GCF, enabling identification of ambient RNAs (containing CD1 SNPs) in GCs purified by FSPS. **(h)** Correlation of genes containing CD1 SNPs with relative mass of CD1 tissue in mixed-strain GCF. **(i)** GO analysis of non-ambient GC RNAs (lacking CD1 SNPs). **(j)** Diluting and centrifuging GCFs pellets GCs, enabling generation of wGCFs via supernatant removal and resuspension. **(k, l)** Representative images **(k)** and quantification **(l)** of western blots against Gap43 and Map2 proteins for GCFs and wGCFs. **(m, n)** RNA **(m)** and protein **(n)** protection assays for wGCFs. Error bars: 95% CI. Statistics: one-sided Student’s t-test (*: p < 0.05; ns: p > 0.05).

**Supplemental Figure 9:**
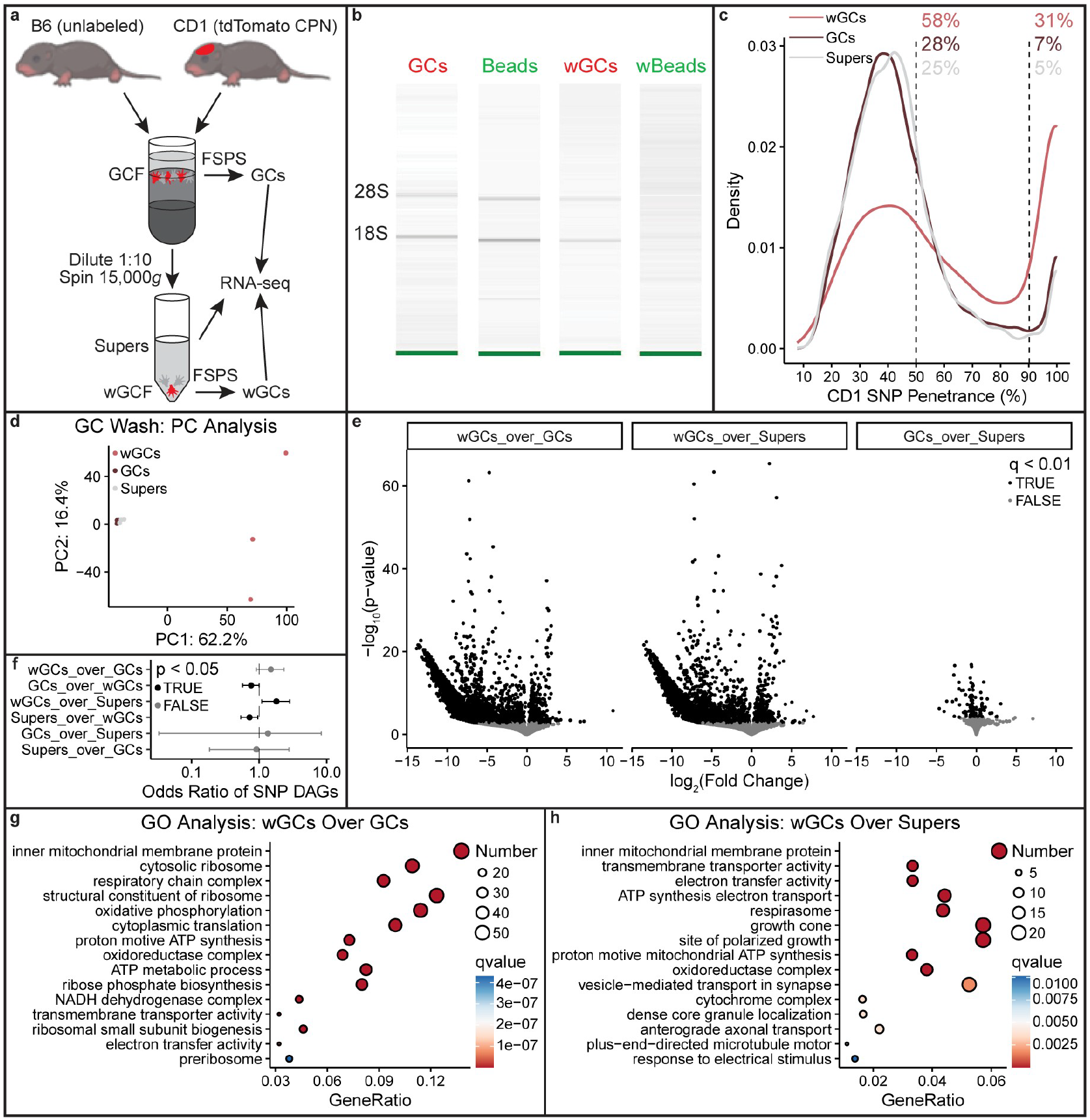
Removal of ambient RNAs from purified CPN GCs. **(a)** Homogenizing unlabeled B6 brains with CD1 brains containing labeled CPN via E14.5 creates a mixed-strain GCF, enabling identification of ambient RNAs (lacking CD1 SNPs) in supernatants, as well as wGCs and GCs, which are purified from wGCFs and GCFs via FSPS, respectively. **(b)** RNA extracted from GCs and Beads, or wGCs and wBeads, which are purified via FSPS from GCFs or wGCFs, respectively. **(c)** CD1 SNP penetrance across samples. **(d)** PC analysis of transcriptomes. **(e)** RNAs with differential abundance across samples. **(f)** Odds ratios of differentially abundant RNAs being enriched for CD1 SNPs. **(g, h)** GO analysis of RNAs enriched in wGCs relative to GCs **(g)** or supers **(h)**. q < 0.1 for volcano plots. Error bars: 95% CI. Odds ratios: two-sided Fisher’s exact test.

## Notes

https://doi.org/10.7910/DVN/KZ2JDL

https://github.com/detillman/lrrtm2_macklis_2026

### Competing Interest Statement

The authors have declared no competing interest.

### Summary of Updates

Updated author affiliations and changed two words in abstract.

